# Long-range allosteric effects - How adenine nucleotides shape the conformational landscape of the Ski2-like RNA helicase, Brr2

**DOI:** 10.1101/2021.01.28.428616

**Authors:** Eva Absmeier, Karen Vester, Tahereh Ghane, Dmitry Burakovskiy, Pohl Milon, Petra Imhof, Marina V. Rodnina, Karine F. Santos, Markus C. Wahl

## Abstract

Brr2 is an essential Ski2-like RNA helicase that exhibits a unique structure among the spliceosomal helicases. Brr2 harbors a catalytically active N-terminal helicase cassette and a structurally similar, but enzymatically inactive C-terminal helicase cassette. Both cassettes contain a nucleotide binding pocket. Here we use biophysical and computational methods to delineate the functional connectivity between the cassettes and how occupancy of the nucleotide binding sites may influence each other. Our results show that Brr2 exhibits high specificity for adenine nucleotides with both cassettes binding ADP tighter than ATP. Adenine nucleotide affinity for the inactive C-terminal cassette is more than two orders of magnitude higher than that of the active N-terminal cassette, largely determined by slow nucleotide release. Mutations at the inter-cassette surfaces and in the connecting linker diminish the affinity of adenine nucleotides for both cassettes. Abrogation of nucleotide binding at the C-terminal cassette reduces nucleotide binding at the N-terminal cassette, 70 Å away. Molecular dynamics simulations identified structural communication lines that likely mediate the long-range allosteric effects. Together, our results reveal intricate networks of intra-molecular interactions in the complex Brr2 RNA helicase, which fine-tune its nucleotide affinities and which could be exploited for regulating the enzyme during splicing.

## INTRODUCTION

Ski2-like nucleic acid helicases constitute a family of superfamily 2 (SF2) helicases. All Ski2-like helicases display a common helicase core consisting of two RecA-like domains, a winged-helix (WH) domain and a helical bundle (HB or “ratchet”) domain arranged in a ring-like fashion (1). This domain arrangement has also been observed in structures of the DEAH/RHA family of SF2 helicases (2-4). Additionally, Ski2-like helicases can contain diverse accessory domains appended to or inserted into the conserved domains, which modulate or expand their molecular mechanisms or functions (5).

The two RecA-like domains contain twelve conserved sequence motifs (“helicase motifs”) involved in nucleotide tri-phosphate (NTP) and nucleic acid transactions (6). Nucleic acid substrates bind with 3’-to-5’ directionality across the first and second RecA-like domains and below the HB/ratchet domain, explaining the 3’-to-5’ unwinding directionality of these enzymes (7-9). Ski2-like helicases also contain a β-hairpin in the second RecA-like domain, which is shorter compared to a similar element in DEAH/RHA helicases, and which is thought to act as a strand separation device that inserts between the two strands of a duplex when one strand of the substrate is pulled through the helicase unit (8,10). Like DEAD-box helicases, Ski2-like helicases are thought to selectively utilize ATP due to the presence of a Q-motif (11).

Four Ski2-like RNA helicases have been identified in yeast: Ski2 (involved in RNA degradation and viral defense) (12), Slh1 (ribosome quality control) (13), Mtr4 (RNA degradation) (7) and Brr2 (pre-mRNA splicing) (14). In addition, an Mtr4 homolog, FRH, of *Neurospora crassa* is involved in the regulation of the circadian rhythm (15). Several additional Ski2-like enzymes are DNA helicases such as yeast Mer3/mammalian HFM1, involved in homologous recombination (16,17), archaeal Hel308/eukaryotic POLQ-like helicase, involved in double strand break repair (8,18), and the mammalian ASCC3 helicase, associated with transcription regulation (19), DNA repair (20), ribosome quality control (21-23), non-functional ribosomal RNA decay (24) and viral defense (25). 3’-to-5’ translocation has been directly shown for Hel308 (8), Mer3/HFM1 (26), Mtr4 (27) and Brr2 (28,29).

A common feature among Ski2-like RNA helicases is their integration into multi-protein assemblies. Brr2 is an integral component of the spliceosomal U5 snRNP (30,31). Thus, during assembly of a spliceosome on a pre-mRNA substrate, Brr2 is recruited as part of the pre-assembled U4/U6•U5 tri-snRNP during formation of a pre-catalytic B complex spliceosome (32-34). Brr2 is essential for the conversion of the pre-catalytic to an activated spliceosome (35,36), during which it unwinds the extensively base-paired U4/U6 di-snRNA (14,37,38). To this end, the enzyme binds to a single-stranded (ss) region of U4 snRNA and translocates in 3’-to-5’ direction (28,38). Unlike other spliceosomal helicases, Brr2 remains stably associated with the spliceosome after incorporation of the tri-snRNP throughout the remaining stages of a splicing event (32,34,39-44), and is required again during the two catalytic steps of splicing (45,46) and during spliceosome disassembly (47). However, the Brr2 helicase activity *per se* may not be necessary during these later stages of splicing (43,45). Thus, Brr2 has to be strictly regulated to prevent premature unwinding of U4/U6 in the tri-snRNP and to allow its repeated on- and off-switching during splicing.

Brr2 is a particularly large spliceosomal RNA helicase (ca. 245 kDa for the human enzyme). The helicase core of Brr2 is expanded by additional helix-loop-helix (HLH) and immunoglobulin-like (IG) domains, which form a Sec63-homology unit together with the HB/ratchet domain (48,49). In addition, Brr2 contains two copies of the RecA1/2-WH-Sec63 units (cassettes) arranged in tandem (Fig. 1A) (50). This dual-cassette organization is shared by only few other Ski2-like enzymes, including the RNA helicase Slh1 (51) and the DNA helicase ASCC3 (20). Only the N-terminal cassette (NC) of Brr2 is an active ATPase and RNA helicase (50) and the enzymatic activity of the NC alone is required for splicing *in vivo* (52). Nevertheless, the C-terminal cassette (CC) is essential for yeast viability (49), represents a versatile protein-protein interaction platform (53,54), retains nucleotide binding capacity (50,55) and regulates the activity of the NC (50). Therefore, the CC may be considered an intra-molecular helicase cofactor in Brr2.

**Figure 1.**
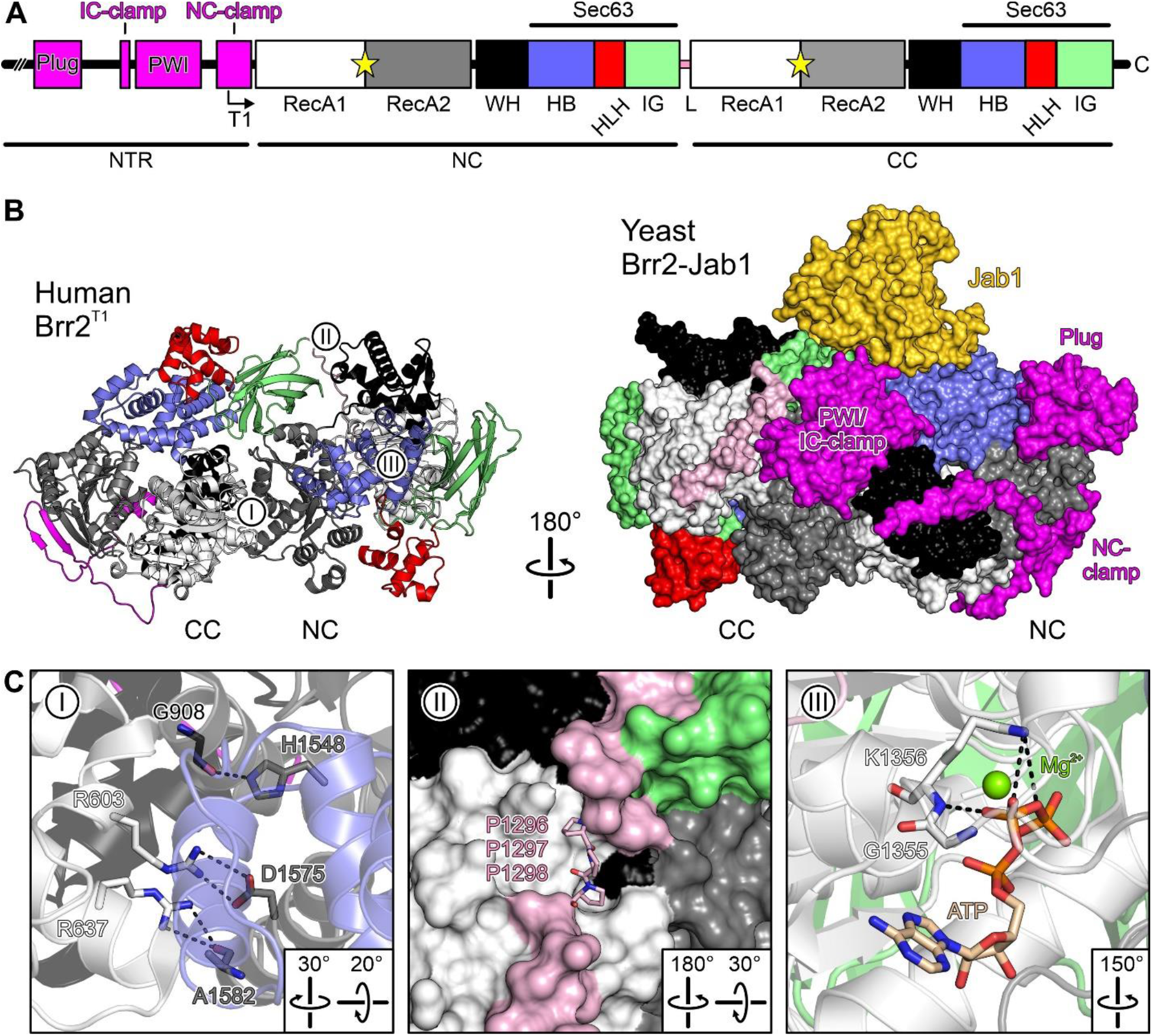
Brr2 organization. **A.** Domain organization of hBrr2. Angled arrow, starting position of the T1 truncation variant. NTR, N-terminal region; NC, N-terminal cassette; CC, C-terminal cassette; RecA, RecA-like domain; WH, winged helix domain; HB, helical bundle domain; HLH, helix-loop-helix domain; IG, immunoglobulin-like domain; L, linker. Yellow stars, nucleotide binding pockets. **B.** Structures of hBrr2^T1^ (PDB ID 4F91; left) and yBrr2^FL^-Jab1 complex (PDB ID 5DCA; right). I-III, regions mutated (see **C**). Domain coloring as in **A**; Jab1, gold. Rotation symbol, relative orientation of the panels. **C.** Details of the mutated regions (I-III as in **A**, left). ATP-bound structure according to PDB ID 4F93 (region III, right). Relevant residues are shown as sticks. Domain coloring as in **A**. Residue coloring: Protein carbon, as the respective protein region; ATP carbon, beige; nitrogen, blue; oxygen, red; phosphorus, orange; magnesium ion, green. Dashed lines, hydrogen bonds or salt bridges.

Preceding its two helicase cassettes, Brr2 contains an evolutionarily conserved N-terminal region (NTR) of approximately 400 residues (Fig. 1A), capable of auto-inhibiting Brr2 (56). The NTR contains two folded domains connected by flexible regions and can fold back along one flank of the tandem helicase cassettes (Fig. 1B), blocking access of the RNA substrate and restricting conformational rearrangements required for RNA duplex unwinding (56). *In vivo*, the NTR is required for yeast viability, efficient splicing, Brr2 association with the U4/U6•U5 tri-snRNP, tri-snRNP stability (56,57), retention of U5 and U6 snRNAs during spliceosome activation (36) and as a protein-protein interaction platform (58).

Mechanistic insights into nucleotide binding, hydrolysis and product release are necessary to understand how NTP transactions are coupled to RNA rearrangements in RNA helicases. In spite of detailed structural analyses of the Brr2 helicase (50,55,56,59), there is limited information about the molecular mechanisms underlying its NTPase cycle. Here, we used biophysical and computational methods to systematically characterize nucleotide binding to wild type (wt) Brr2 and selected variants. Our results show that the Brr2 helicase cassettes have drastically different affinities to nucleotides and that ADP is preferred over ATP in both nucleotide binding pockets. Residue exchanges at the inter-cassette interface reduced nucleotide association for both cassettes and mutations in the CC nucleotide-binding pocket interfered with nucleotide binding at the NC. We delineated structural communication lines in the enzyme by molecular dynamics simulations, which may mediate these long-range effects. Our results show that accessory regions in an RNA helicase (CC in the case of Brr2) can regulate nucleotide affinity and specificity in an intricate manner.

## RESULTS

### Nucleotide specificity of human Brr2

We used recombinant human Brr2 (hBrr2), which we consider a good representative of all Brr2 orthologs, as Brr2 is evolutionarily highly conserved. Apart from full-length hBrr2, we investigated an N-terminally truncated variant (T1) that lacks most of the auto-inhibitory N-terminal region as well as the isolated helicase cassettes (hBrr2^NC^, hBrr2^CC^; Fig. 1A).

hBrr2 has two nucleotide binding pockets, one in each of its cassettes (50) (Fig. 1A). The CC pocket is inactive in ATP hydrolysis (50). We first set out to directly test nucleotide binding preferences for each cassette of hBrr2 in isolation, hBrr2^NC^ (residues 395-1324) and hBrr2^CC^ (residues 1282-2136), making use of fluorescence resonance energy transfer (FRET) from tryptophan residues in the vicinity of the NC and CC nucleotide binding pockets (Trp817 in hBrr2^NC^ or Trp1652 and Trp1393 in hBrr2^CC^) to *mant*-nucleotides. In our experimental setups, excitation of tryptophans resulted in *mant* fluorescence increases upon binding of the labeled nucleotides; conversely, dissociation of the *mant*-nucleotides resulted in decreased FRET. Time courses of binding were recorded at a constant concentration of hBrr2^NC^ or hBrr2^CC^ (0.5 μM) with an excess of *mant*-nucleotide (5 μM; Fig. 2). Among the *mant*-nucleotides tested (ADP, ATP, ATPγS, AMPPNP, GDP, GTP, GTPγS), only *mant*-ADP and *mant-* ATPγS showed FRET with both hBrr2^NC^ and hBrr2^CC^ (Fig. 2A,B). *mant*-ATP only yielded a FRET signal with hBrr2^CC^, but not with hBrr2^NC^ (Fig. 2A,B). None of the guanosine nucleotides showed binding to either cassette.

**Fig. 2.**
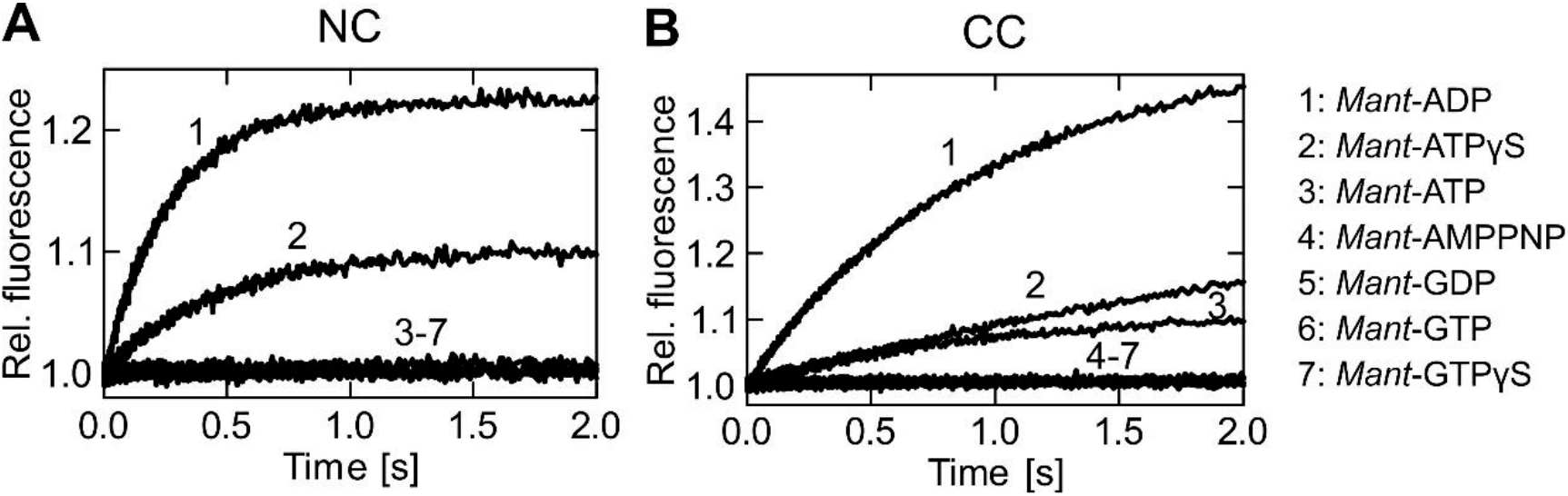
Nucleotide specificity of the hBrr2 cassettes. **A** and **B**. Time courses of *mant*-nucleotide binding to 0.5 μM nucleotide-free hBrr2^NC^ (**A**) and hBrr2^CC^ (**B**) measured by FRET from Trp to *mant.* 1, *mant*-ADP (5 μM); 2, *mant*-ATPγS (5 μM); 3, *mant*-ATP (5 μM); 4, *mant*-AMPPNP (5 μM); 5, *mant*-GDP (5 μM); 6, *mant*-GTP (5 μM); 7, *mant*-GTPγS (5 μM). The hBrr2 cassettes bind *mant*-ADP and *mant*-ATPγS but do not interact with *mant*-AMPPNP or *mant*-G nucleotides.

These data show that both cassettes of hBrr2 possess a high specificity for adenine nucleotides, as expected from the presence of Q-motifs in the RecA1 domains of both cassettes, and suggested that both preferentially bind ADP over ATPγS.

### Structures of mant-nucleotides bound at NC and CC

Lack of a FRET signal from *mant*-AMPPNP with either cassette (Fig. 2A,B) may indicate that AMPPNP is not a suitable ATP surrogate in the case of hBrr2. Similar observations have been made with GMPPNP and some GTPases (60). However, lack of a FRET signal from *mant*-ATP with hBrr2^NC^ was surprising, given that *mant*-ATPγS yielded a signal (Fig. 2A,B). Based on this observation, we set out to investigate whether *mant*-nucleotides that give rise to FRET signals exhibit the same binding poses as the corresponding unlabeled nucleotides, and whether the *mant* moieties influence nucleotide binding. To this end, we determined crystal structures of hBrr2 in complex with ADP, ATPγS, *mant*-ADP and *mant*-ATPγS at 2.5-2.8 Å resolution (Table S1). For crystallographic analyses, we used N-terminally truncated hBrr2^T1^ in complex with the Jab1 domain of the hPrp8 protein lacking a Brr2-inhibiting C-terminal tail (hJab1^ΔC^). The hBrr2^T1^-hJab1^ΔC^ complex yields well-diffracting crystals under low-salt conditions (61), suitable for nucleotide binding.

Electron densities for the phosphate groups, the riboses and the adenine bases were well defined in both NC and CC for all nucleotides analyzed (Fig. 3A-D). At both cassettes, accommodation of the phosphate/ribose/base portions of *mant-ADP* and *mant-ATPγS* were essentially unaltered compared to ADP and ATPγS (Fig. 3A-D), showing that the *mant* units do not influence how the nucleotides are positioned in the binding pockets. Moreover, ATPγS and the ATPγS portion of *mant*-ATPγS are accommodated as expected for ATP (50). The structures also revealed that two tryptophan residues (W1393 and W1652) reside within 20 Å distance of the CC nucleotide binding pocket, while only one tryptophan residue (W817) is close to the NC nucleotide binding pocket (Fig. S1), providing an explanation for higher FRET at the CC.

**Figure 3.**
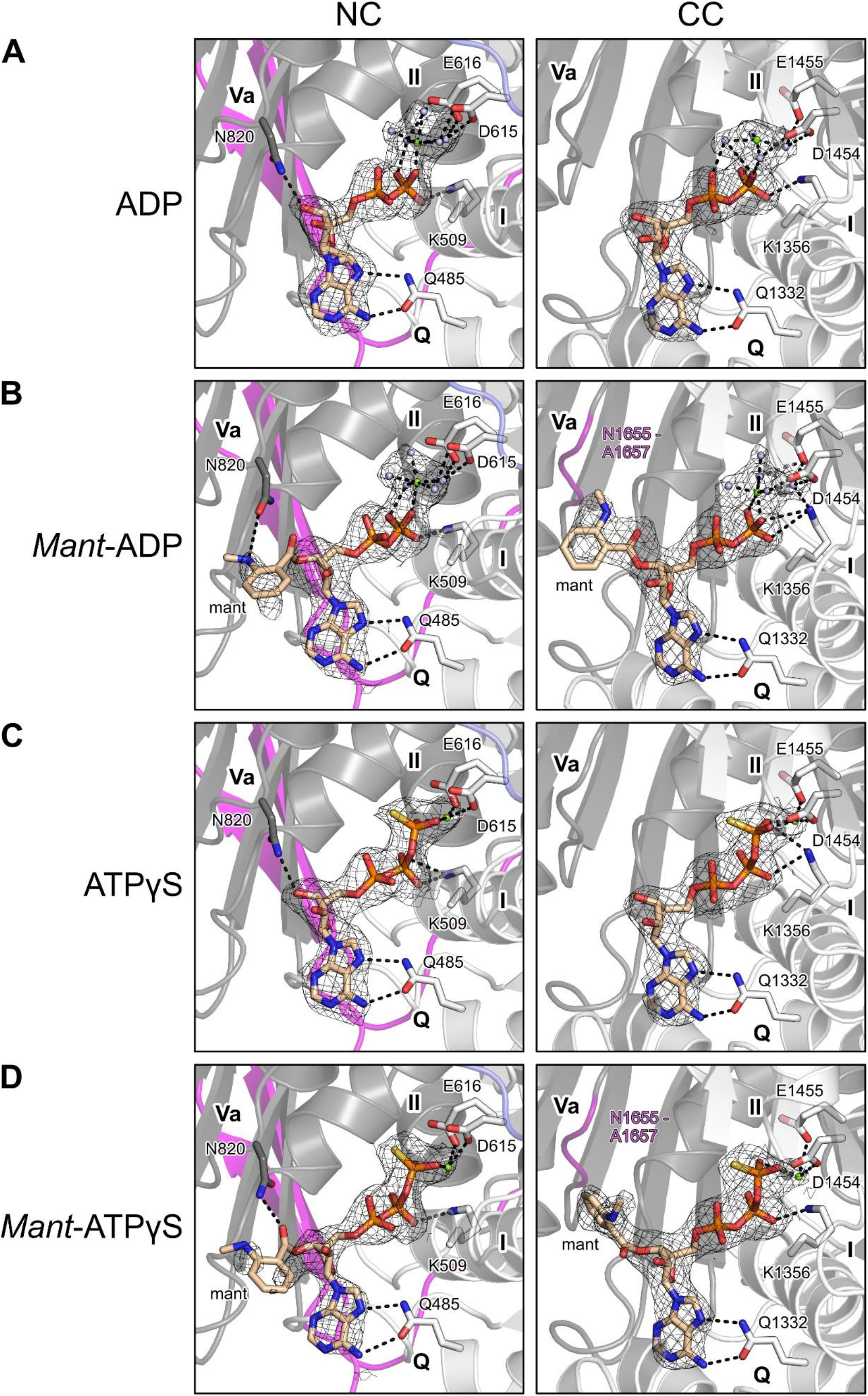
Nucleotide-bound structures. **A-D.** Binding poses of the indicated nucleotides at the N- and C-terminal cassettes of a hBrr2^T1^-hJab1^ΔC^ complex. Meshes, final 2mF_o_-DF_c_ electron density maps at the 1σ level coving the nucleotides. Left panels, nucleotides bound at NC; right panels, nucleotides bound at CC. Nucleotides, magnesium ions, selected waters and selected Brr2 residues are shown as sticks or small spheres. Domain coloring as in Fig. 1A. Atom/ion coloring: Protein carbon, as the respective protein region; nucleotide carbon, beige; nitrogen, blue; oxygen, red; phosphorus, orange; magnesium ion, green; water oxygen, light blue. Dashed lines, hydrogen bonds, salt bridges or ion coordination.

Additionally, electron densities for the *mant* units of *mant*-ADP and *mant*-ATPγS were less defined in the NC compared to the CC (Fig. 3A-D), indicating that the fluorophores remain more mobile when *mant*-nucleotides are accommodated at the NC compared to the CC. With *mant* nucleotides bound at the NC, N820 (helicase motif Va) fosters a hydrogen bond to the *mant* moieties instead of the sugar units of unlabeled nucleotides (Fig. 3A-D, left). At the CC, *mant* moieties engage in additional van der Waals (vdW) contacts to the helicase motif Va residues (Fig. 3B,D, right).

Based on these results, we conclude that the *mant* moieties do not significantly alter the positioning of ADP or ATPγS in the nucleotide binding pockets, but may lead to higher affinities of *mant*-nucleotides compared to the corresponding unmodified nucleotides by fostering additional contacts. The better defined electron densities for the *mant* moieties in the CC compared to the NC suggest that the *mant* units have a relatively larger effect on nucleotide affinities to the CC compared to the NC. Furthermore, comparable distances (within ± 0.2 Å) between *mant* moieties and close Trp residues in the structures with *mant*-ADP and *mant*-ATPγS (Fig. S1), suggest that higher FRET signals obtained with *mant*-ADP compared to *mant*-ATPγS (Fig. 2A,B) are due to a lower occupancy of the nucleotide binding pockets in the case of *mant-ATPγS* under the chosen conditions, as a consequence of a lower affinity of ATPγS compared to ADP. In addition, we suggest that hBrr2^NC^ exhibits higher conformational flexibility than the NC in context of a dual-cassette Brr2 construct (hBrr2^T1^ or hBrr2^FL^), and that, therefore, *mant*-ATP is not sufficiently stably bound at the isolated NC to yield a FRET signal (Fig. 2A).

We were predominantly interested in affinity changes due to allosteric effects, which we expect to affect *mant*-nucleotides and unmodified nucleotides in a similar manner. We, thus, concluded that *mant*-ADP and *mant*-ATPγS are suitable surrogates for studying the binding of ADP and ATP to the Brr2 nucleotide binding pockets and, therefore, employed these nucleotides in subsequent experiments.

### Affinities of the isolated N- and C-terminal cassettes of hBrr2 for adenine nucleotides

Our initial investigations on the nucleotide specificity of the hBrr2 cassettes in isolation indicated that hBrr2^NC^ and hBrr2^CC^ bind nucleotides with different rates. To determine the nucleotide association rate constants, *k_1_^NC^* and *k_1_^CC^*, and the equilibrium affinity constants, we investigated *mant*-ATPγS/*mant*-ADP binding to hBrr2^NC^ and hBrr2^CC^ using strategies previously described for G nucleotide binding to translation factor GTPases (62-65).

We acquired time courses of binding at a constant concentration of nucleotide-free hBrr2^NC^ or hBrr2^CC^ and varying concentrations of *mant*-ATPγS/*mant*-ADP under pseudo-first order conditions (Fig. 4A,B). The time dependencies of FRET were analyzed by exponential fitting to calculate the apparent rate constant of nucleotide binding to either cassette at each nucleotide concentration (*k_app_^NC^* and *k_app_^CC^*). hBrr2^NC^ and hBrr2^CC^ time traces were best fit with a single-exponential equation, consistent with a one-step binding model (Fig. 4A,B). The bimolecular association rate constants, *k_1_^NC^* and *k_1_^CC^*, were determined from the slopes of the linear dependences of *k_app_^NC^* and *k^app^^CC^*, respectively, on the concentration of *mant*-ATPγS/*mant*-ADP (Fig. 4C,D; Table 1). The values obtained indicate that both isolated hBrr2^NC^ and hBrr2^CC^ bind ADP faster than ATPγS. Furthermore, hBrr2^NC^ binds both tested nucleotides faster than hBrr2^CC^.

**Fig. 4.**
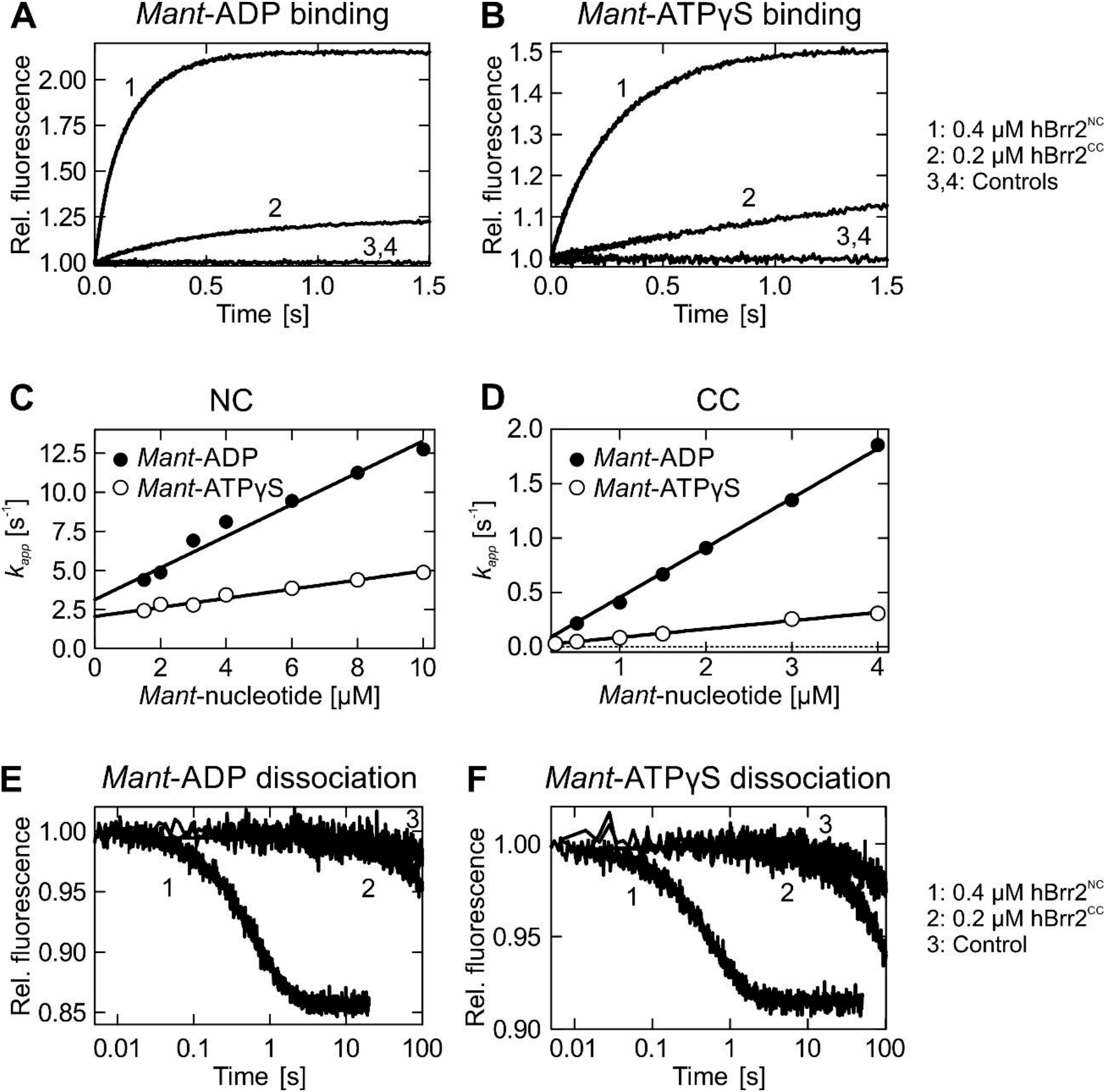
Kinetics of *mant-ADP* and *mant*-ATPγS interaction with hBrr2^NC^ and hBrr2CC. **A** and **B**. Time courses of 5 μM *mant*-ADP (**A**) or *man*t-ATPγS (**B**) binding to hBrr2^NC^ (1; 0.4 μM) and hBrr2CC (2; 0.2 μM) measured by FRET from Trp to *mant*. Controls correspond to binding experiments with unlabeled ADP or ATPγS (3 and 4). **C** and **D**. Individual nucleotide binding traces were fitted to single exponentials and the dependencies of the apparent rate constants on nucleotide concentration for hBrr2^NC^ (**C**) and hBrr2CC (**D**) were fitted by a linear equation, *k_app_* = *k_1_*[mant-nucleotide]+*k-_1_*, in which *k_1_* is derived from the slope and *k-_1_* from the Y-axis intercept. Closed circles, *mant*-ADP; open circles, *mant*-ATPγS. Values represent means ± SD of at least three independent measurements. **E** and **F**. Dissociation of 5 μM *mant*-ADP (**E**) or *mant*-ATPγS (**F**) from hBrr2^NC^ (1; 0.4 μM hBrr2^NC^) and hBrr2^CC^ (2; 0.2 μM) in the presence of the respective unlabeled nucleotide in excess (100 μM). Control experiments (3) were carried out in the absence of unlabeled nucleotide (curves shown are for hBrr2^CC^).

**Table 1.** Kinetics of nucleotide binding and release.

| Protein | Data fitting<sup>a</sup> | *mant*-ADP | | | | *mant*-ATPγS | | | |
| --- | --- | --- | --- | --- | --- | --- | --- | --- | --- |
|  |  | NC |  | CC |  | NC |  | CC |  |
| | | $k_1$<br>[$\mu\text{M}^{-1} \text{s}^{-1}$] | $k_{-1}$<br>[$\text{s}^{-1}$] | $k_1$<br>[$\mu\text{M}^{-1} \text{s}^{-1}$] | $k_{-1}$<br>$10^{-3} [\text{s}^{-1}]$ | $k_1$<br>[$\mu\text{M}^{-1} \text{s}^{-1}$] | $k_{-1}$<br>[$\text{s}^{-1}$] | $k_1$<br>[$\mu\text{M}^{-1} \text{s}^{-1}$] | $k_{-1}$ $10^{-3}$<br>[$\text{s}^{-1}$] |
| **hBrr2<sup>NC</sup>** | SE | 1.0<br>$\pm 0.1$ | 1.9<br>$\pm 0.1$ | - | - | 0.3<br>$\pm 0.02$ | 1.6<br>$\pm 0.02$ | - | - |
| **hBrr2<sup>CC</sup>** | SE | - | - | 0.5<br>$\pm 0.01$ | 2.0<br>$\pm 0.2$ | - | - | 0.1<br>$\pm 0.001$ | 2.0<br>$\pm 0.2$ |
| **hBrr2<sup>FL</sup>** | DE | 2.8<br>$\pm 0.03$ | 1.5<br>$\pm 0.02$ | 0.4<br>$\pm 0.01$ | 1.0<br>$\pm 0.1$ | 1.0<br>$\pm 0.02$ | 1.5<br>$\pm 0.04$ | 0.1<br>$\pm 0.01$ | 1.0<br>$\pm 0.1$ |
| **hBrr2<sup>T1</sup>** | DE | 2.5<br>$\pm 0.1$ | 1.3<br>$\pm 0.02$ | 0.4<br>$\pm 0.02$ | 1.0<br>$\pm 0.1$ | 2.5<br>$\pm 0.03$ | 1.2<br>$\pm 0.04$ | 0.1<br>$\pm 0.01$ | 0.9<br>$\pm 0.01$ |
| **R603A** | DE | 0.2<br>$\pm 0.03$ | 1.4<br>$\pm 0.04$ | 0.02<br>$\pm 0.001$ | 1.0<br>$\pm 0.1$ | 0.3<br>$\pm 0.03$ | 1.0<br>$\pm 0.04$ | 0.02<br>$\pm 0.001$ | 1.0<br>$\pm 0.2$ |
| **R637A** | DE | 3.4<br>$\pm 0.2$ | 1.2<br>$\pm 0.03$ | 0.5<br>$\pm 0.01$ | 1.0<br>$\pm 0.1$ | 0.4<br>$\pm 0.04$ | 1.0<br>$\pm 0.1$ | 0.02<br>$\pm 0.001$ | 1.0<br>$\pm 0.04$ |
| **S1087L<sup>b</sup>** | DE | 2.7<br>$\pm 0.1$ | 1.4<br>$\pm 0.04$ | 0.5<br>$\pm 0.01$ | 1.0<br>$\pm 0.03$ | 2.2<br>$\pm 0.21$ | 1.4<br>$\pm 0.04$ | 0.1<br>$\pm 0.004$ | 1.0<br>$\pm 0.4$ |
| **PPP1296-8AAA** | DE | 2.9<br>$\pm 0.1$ | 2.2<br>$\pm 0.1$ | 0.52<br>$\pm 0.003$ | 2.0<br>$\pm 0.04$ | 0.5<br>$\pm 0.03$ | 2.2<br>$\pm 0.1$ | 0.03<br>$\pm 0.002$ | 2.0<br>$\pm 0.4$ |
| **GK1355-6QE** | SE | 1.3<br>$\pm 0.1$ | 1.5<br>$\pm 0.02$ | - | - | 0.4<br>$\pm 0.02$ | 1.7<br>$\pm 0.1$ | - | - |
| **H1548A** | DE | 3.3<br>$\pm 0.3$ | 1.2<br>$\pm 0.1$ | 0.5<br>$\pm 0.01$ | 1.0<br>$\pm 0.3$ | 0.2<br>$\pm 0.02$ | 1.3<br>$\pm 0.1$ | 0.02<br>$\pm 0.001$ | 1.0<br>$\pm 0.03$ |

<sup>a</sup> SE, single-exponential; DE, double exponential.<sup>b</sup> S1087L in the N-terminal HB domain originates from a retinitis pigmentosa-linked mutation of hBrr2, and was carried along as a negative control.**Table 2.** Affinities of hBrr2 variants for *mant*-ADP and *mant*-ATPγS.

The dissociation rate constants were determined by pre-forming hBrr2^NC/CC^ complexes with *mant*-ADP or *mant*-ATPγS, and mixing the complexes with an excess of the respective unlabeled nucleotide, using a stopped-flow apparatus. Under these conditions, the rate of ADP/ATPγS binding is limited by the rate of *mant*-ADP/*mant*-ATPγS dissociation from hBrr2^NC^ or hBrr2^CC^. Rebinding of the *mant*-nucleotide is negligible due to the large excess of unlabeled nucleotide. Thus, the rate by which the *mant* fluorescence decreases equals the dissociation rate of the *mant*-nucleotide. Dissociation rate constants were obtained by single-exponential fitting of the time courses (Fig. 4E,F; Table 1). hBrr2^NC^ dissociation rates (*k-_1_^NC^*) were 1.9±0.1 s^-1^ and 1.6±0.02 for *mant*-ADP and *mant*-ATPγS, respectively. ADP and ATPγS dissociation rates for hBrr2^CC^ (*k-_1_^CC^*) were three orders of magnitude lower than for hBrr2^NC^, 2 ± 0.2 10-^3^ s^-1^ (Fig. 4E,F; Table 1).

The *K_d_* values for the interaction of ADP and ATPγS with hBrr2^NC^ and hBrr2^CC^ were calculated from the ratios of dissociation and association rate constants (*K_d_^NC^* = *k-_1_^NC^/k_1_^NC^; K_d_^CC^* = *k-_1_^CC^*/*k_1_^CC^*). Both hBrr2^NC^ (3-fold) and hBrr2^CC^ (7.5-fold) displayed a higher affinity toward ADP compared to ATPγS (Table 2). Furthermore, hBrr2^CC^ has about 500-fold and about 200-fold higher affinities for ADP and ATPγS, respectively, as compared to hBrr2^NC^, largely determined by the dissociation rates (*k-_1_^CC^*) for *mant*-nucleotides (Table 2). Thus, although the two helicase cassettes are structurally similar, they differ over more than two orders of magnitude in their capacity to interact with adenine nucleotides.

**Table 2.** Affinities of hBrr2 variants for *mant*-ADP and *mant*-ATPγS.

| Protein | *mant*-ADP | | *mant*-ATPγS | |
| --- | --- | --- | --- | --- |
|  | NC | CC | NC | CC |
| | $K_d$<br>[$\mu\text{M}$] | $K_d$<br>[nM] | $K_d$<br>[$\mu\text{M}$] | $K_d$<br>[nM] |
| **hBrr2<sup>NC</sup>** | 2.0±0.2 | - | 5.8±0.4 | - |
| **hBrr2<sup>CC</sup>** | - | 4±0.4 | - | 30±3 |
| **hBrr2<sup>FL</sup>** | 0.5±0.01 | 2±0.2 | 1.6±0.1 | 10±1 |
| **hBrr2<sup>T1</sup>** | 0.5±0.02 | 2±0.3 | 0.5±0.02 | 10±1 |
| **R603A** | 6.3±0.8 | 50±4 | 4.1±0.5 | 50±10 |
| **R637A** | 0.4±0.2 | 2±0.1 | 2.6±0.03 | 40±2 |
| **S1087L<sup>a</sup>** | 0.5±0.02 | 2±0.1 | 0.6±0.1 | 10±4 |
| **PPP1296-8AAA** | 0.8±0.05 | 4±0.1 | 4.3±0.4 | 60±10 |
| **GK1355-6QE** | 1.2±0.05 | - | 4.4±0.3 | - |
| **H1548A** | 0.4±0.04 | 2±0.6 | 5.9±0.6 | 40±2 |

<sup>a</sup> S1087L in the N-terminal HB domain originates from a retinitis pigmentosa-linked mutation of hBrr2, and was carried along as a negative control.

### Interaction of mant-ATPγS/mant-ADP with hBrr2^FL^ and hBrr2^T1^

We next studied nucleotide binding behavior to hBrr2^FL^ and hBrr2^T1^, using the same detection principles as described above. hBrr2^FL^ comprises the full-length helicase, whereas hBrr2^T1^ lacks the NTR except for a segment that meanders along the NC (NC-clamp; Fig. 1A). The time courses of nucleotide binding were obtained at a constant concentration of nucleotide-free hBrr2^FL^ or hBrr2^T1^ and a varying concentration of *mant-*ADP/*mant*-ATPγS. For both, hBrr2^FL^ and hBrr2^T1^, biphasic time dependencies were observed (Fig. 5A,B), consistent with the nucleotides binding to the NC and CC of hBrr2. Nonlinear regression analysis using two exponential terms yielded two apparent rate constants, *k_app1_* and *k_app2_*, for each time trace. To assign each *k_app_* to NC or CC, we integrated the results as for the isolated cassette constructs. *k_app1_* dependencies over nucleotide concentration were in the same range as those obtained with the isolated hBrr2^NC^, whereas *k_app2_* dependencies roughly agreed with those of hBrr2^CC^, allowing assignment of each apparent rate constant to adenine nucleotide interaction with each cassette. The bimolecular association rate constants, *k_1_^NC^* and *k_1_^CC^*, were determined from the linear concentration dependences of *k_app_^NC^* and *k_app_^CC^* on the concentration of *mant*-nucleotides (Fig. 5C,D; Table 1).

**Fig. 5.**
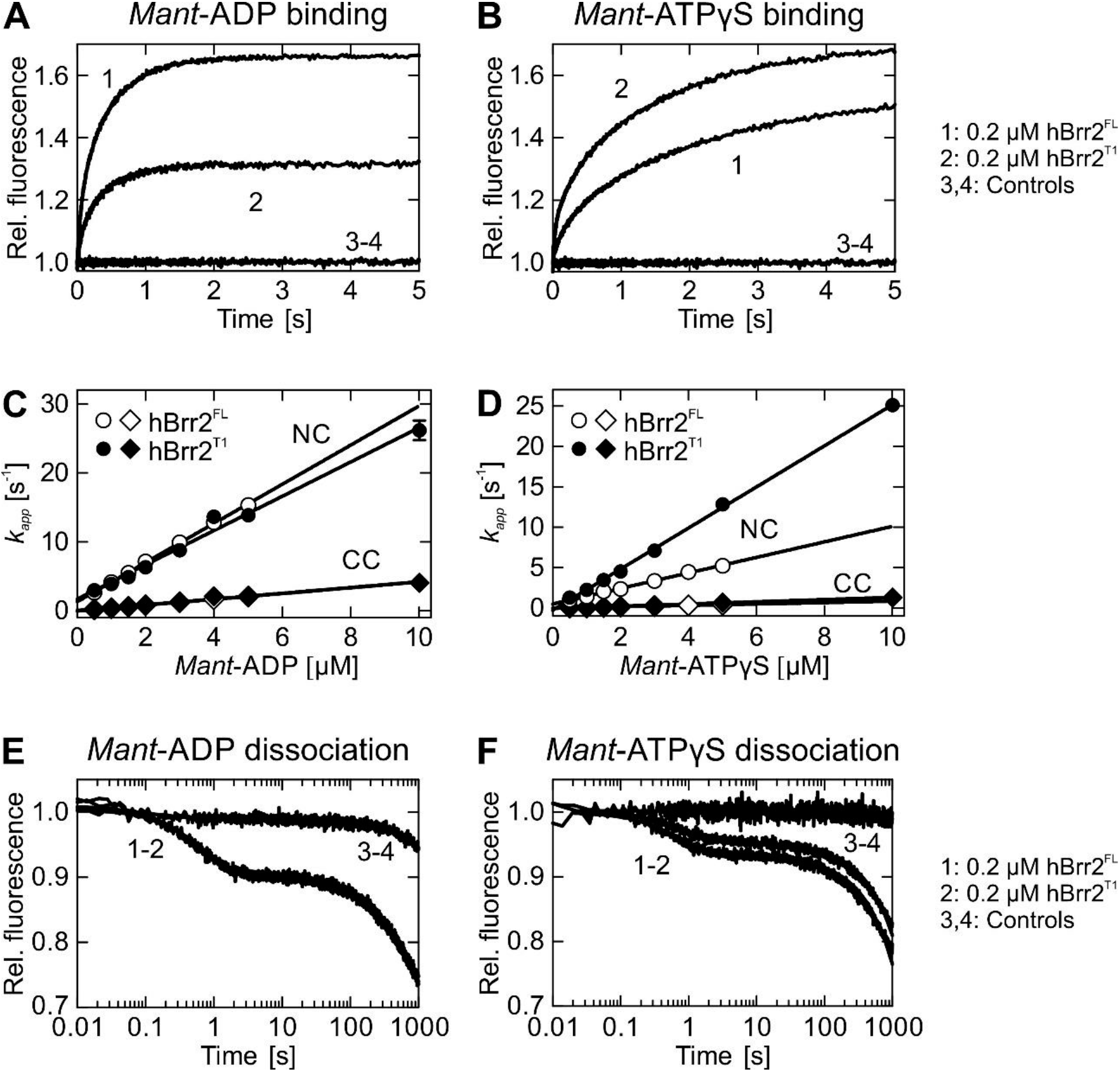
Kinetics of *mant-ADP* and *mant*-ATPγS interaction with hBrr2^FL^ and hBrr2^T1^. **A** and **B**. Time courses of 5 μM *mant*-ADP (**A**) or *man*t-ATPγS (**B**) binding to hBrr2^FL^ (1; 0.2 μM) and hBrr2^T1^ (2; 0.2 μM) measured by FRET from Trp to *mant*. Controls were performed with unlabeled ADP or ATPγS (3-4). **C** and **D**. Individual nucleotide binding traces were fitted to double exponential equations, and the dependence of the apparent rate constants of the NC (circles) and CC (diamonds) on nucleotide concentration were fitted by a linear equation. Open symbols, hBrr2^FL^; closed symbols, hBrr2^T1^. The cassettes of hBrr2^FL^ and hBrr2^T1^ bind nucleotides with different velocities as observed for the variants containing either one of the cassettes, hBrr2^NC^ and hBrr2^CC^. While the CC binds *mant*-ADP and *mant*-ATPγS with similar rates, the NC in hBrr2^T1^ binds *mant*-ATPγS faster than the NC in hBrr2^FL^. Values represent means ± SD of at least three independent measurements. **E** and **F**. Dissociation of 5 μM *mant-ADP* (**E**) or *mant*-ATPγS (**F**) from hBrr2^FL^ (1; 0.2 μM) and hBrr2^T1^ (2; 0.2 μM) in the presence of the respective unlabeled nucleotide in excess (100 μM). Control experiments (3-4) were carried out in the absence of unlabeled nucleotide.

The slope of the linear fitting of *k_app_^NC^*, corresponding to the binding of *mant*-ADP to the NC in the context of hBrr2^FL^ and hBrr2^T1^, indicated association rate constants *k_1_^NC^* of 2.8±0.03 and 2.5±0.11 μM^-1^ s^-1^, respectively. The *k_1_^NC^* values obtained for *mant*-ATPγS binding to the NC of hBrr2^FL^ and hBrr2^T1^ were 1.0±0.02 and 2.5±0.03 μM^-1^ s^-1^, respectively. The *mant*-ADP and *mant*-ATPγS association rates to the NC in the context of hBrr2^FL^ and hBrr2^T1^ are thus ranging from 8-to 3-fold faster compared to the rates of nucleotide binding to isolated hBrr2^NC^ (*k_1_^NC^* of 1.0±0.07 μM^-1^ s^-1^ and 0.3±0.02, respectively, for *mant*-ADP and *mant*-ATPγS), suggesting that the CC slightly improves the ability of the NC to bind nucleotides. These observations are in line with previous findings that hBrr2^NC^ exhibits lower intrinsic and stimulated ATPase activities and lower U4/U6 di-snRNA unwinding activity compared to hBrr2^T1^ (50).

*Mant*-ADP binding to the CC of hBrr2^FL^ and hBrr2^T1^ was faster compared to *mant*-ATPγS, with an association rate constant, *k_1_^CC^*, of 0.4±0.01 μM^-1^ s^-1^ for both hBrr2 variants. The hBrr2^FL^ and hBrr2^T1^ rate constants for *mant*-ATPγS binding to the CC, *k_1_^CC^*, were similar (0.1±0.01 μM^-1^ s^-1^). These values agree well with the *k_1_^CC^* obtained for *mant*-ADP and *mant*-ATPγS binding to the isolated hBrr2^CC^, 0.5±0.01 and 0.1±0.001 μM^-1^ s^-1^, respectively, indicating that presence of the NC does not significantly influence nucleotide binding at the CC.

Nucleotide dissociation from hBrr2^FL^ and hBrr2^T1^ was studied as before with the isolated hBrr2^NC^ and hBrr2^CC^ constructs. Nucleotide dissociation rate constants, *k-_1_^NC^* and *k-_1_^CC^*, were determined upon mixing hBrr2^FL/T1^-*mant*-ADP or hBrr2^FL/T1^-*mant*-ATPγS with an excess of the respective unlabeled nucleotide. The release of the labeled nucleotide from both hBrr2 nucleotide binding pockets resulted in a two-phase fluorescence decrease, consistent with the dissociation of *mant*-ADP or *mant*-ATPγS from the two hBrr2 cassettes (Fig. 5E,F). As nucleotide dissociation from the isolated NC was much faster than from the isolated CC (see above), the dissociation rate constants of the fast phases were assigned to the NC, *k-_1_^NC^*. The nucleotide dissociation rate constants *k-_1_^NC^* for hBrr2^FL^ and hBrr2^T1^ were very similar, around 1.5±0.02 (hBrr2^FL^) and 1.3±0.02 s^-1^ (hBrr2^T1^) for *mant*-ADP and around 1.5±0.04 s^-1^ (hBrr2^FL^) and 1.2±0.04 s^-1^ (hBrr2^T1^) for *mant-*ATPγS (Table 1). These values are in agreement with the *k-_1_^NC^* obtained for *mant-* ADP and *mant*-ATPγS in the isolated hBrr2^NC^ (1.9±0.1 s^-1^ and 1.6±0.02, respectively). The second phase indicated very slow nucleotide release from the CC in the context of hBrr2^FL^ and hBrr2^T1^, with a dissociation rate constant, *k-_1_^CC^*, of about 1 ± 0.1 10^-3^ s^-1^ for ADP and ATPγS, similar to the values associated with the very slow release of nucleotides from isolated hBrr2^CC^. Therefore, the CC appears to be primarily a binding site where nucleotides interact and remain bound due to the low dissociation rate constants. The NC is a site of high turnover with rapid nucleotide binding and rapid release, consistent with its abilities to hydrolyze ATP and promote RNA unwinding (50).

The *K_d_* values for the interaction of ADP and ATPγS with hBrr2^FL^ and hBrr2^T1^ were also calculated from the ratios of the association (*k_1_^NC^* and *k_1_^CC^*) and dissociation rate constants (*k-_1_^NC^* and *k-_1_^CC^*). Due to the small dissociation rate constants, *k-_1_^CC^*, the CC nucleotide binding pockets in hBrr2^FL^/hBrr2^T1^ have about 270-fold/260-fold higher affinities for ADP and about 160-fold/50-fold higher affinities for ATPγS, respectively, compared to the NC pocket (Table 2).

Both nucleotide binding pockets of hBrr2^FL^ exhibit slightly higher affinities for ADP than ATPγS, with a 3-fold and 5-fold preference for ADP in the NC and CC, respectively. Upon NTR truncation (hBrr2^T1^), ADP and ATPγS bind with equal affinities to the NC, while the CC still has a five-fold higher affinity for ADP. Thus, presence of the NTR seems to bias the relative NC nucleotide preference towards ADP, which may contribute to its function as an auto-inhibitory device.

### Residues in the inter-cassette interface and linker modulate nucleotide binding at both cassettes

Direct contacts of the NC RecA1 domain to the CC RecA2 domain observed in Brr2 crystal structures (50,55,56,59) are candidate sites for inter-cassette communication (Fig. 1C, left). Indeed, single alanine substitutions expected to weaken interactions between the NC RecA1 and the CC RecA2 domains have been reported to reduce the helicase activity of hBrr2^T1^ without affecting its RNA binding properties (50). The linker between NC and CC also establishes interaction networks between the cassettes, and alterations to the linker can likewise affect hBrr2^T1^ helicase activity both positively and negatively (50). We therefore investigated if inter-cassette and linker mutants also affect nucleotide binding at either hBrr2 cassette. hBrr2^T1^ mutants were expressed, purified and exhibited cooperative transitions with comparable melting temperatures in thermofluor-based thermal melting experiments (Fig. S2). Furthermore, equilibrium CD spectra were indicative of a high content of regular secondary structure in all hBrr2 variants. These data indicate that all hBrr2^T1^ variants tested herein were well folded, and that mutant effects were not simply a result of a loss of stable tertiary structure.

Rate constants of nucleotide binding and dissociation to hBrr2^T1^ mutants were determined as described above for the wt hBrr2^T1^ protein. All association and dissociation time courses recorded for interface and linker mutants were best described by double-exponential fitting, representing the nucleotide binding and dissociation events at the NC and CC. The bimolecular association rate constants for nucleotide binding at the NC and CC, *k_1_^NC^* and *k_1_^CC^*, were determined from the slope of the linear concentration dependence of the *k_app_^NC^* and *k_app_^CC^* (Table 1). Strikingly, all variants, R603A (NC-RecA1), R637A (NC-RecA1), PPP1296-1298AAA (linker) and H1548A (CC-RecA2; Fig. 1C, left and middle), exhibited reduced ATPγS association rate constants to the NC and CC, while the dissociation rate constants for both cassettes were similar to those of wt hBrr2^T1^ (Fig. 6A,B). As a consequence, the resultant *K_d_*’s showed that the variant proteins bind ATPγS with lower affinity compared to wt hBrr2^T1^. Interestingly, only one mutant had an effect on ADP binding. The R603A variant displayed about 10-fold and about 20-fold lower ADP association rate constants for NC and CC, respectively, compared to wt hBrr2^T1^ (Fig. 6C,D; Table 1). ADP dissociation rate constants of both cassettes were largely unaffected by the mutations (Fig. 6C,D; Table 1). In summary, all mutants reduced binding of ATPγS to either cassette, one mutant (R603A) also reduced ADP binding to both cassettes, while the release of ADP or ATPγS from either cassette was largely unaffected. Thus, inter-cassette contacts appear to configure the adenine nucleotide binding pockets for ATP binding. Interestingly, ATPγS association rate constants to hBrr2^T1^ variants with diminished inter-cassette contacts resemble those of the isolated hBrr2^NC^, suggesting that the regulatory CC requires those inter-cassette contacts to modulate the NC.

**Fig. 6.**
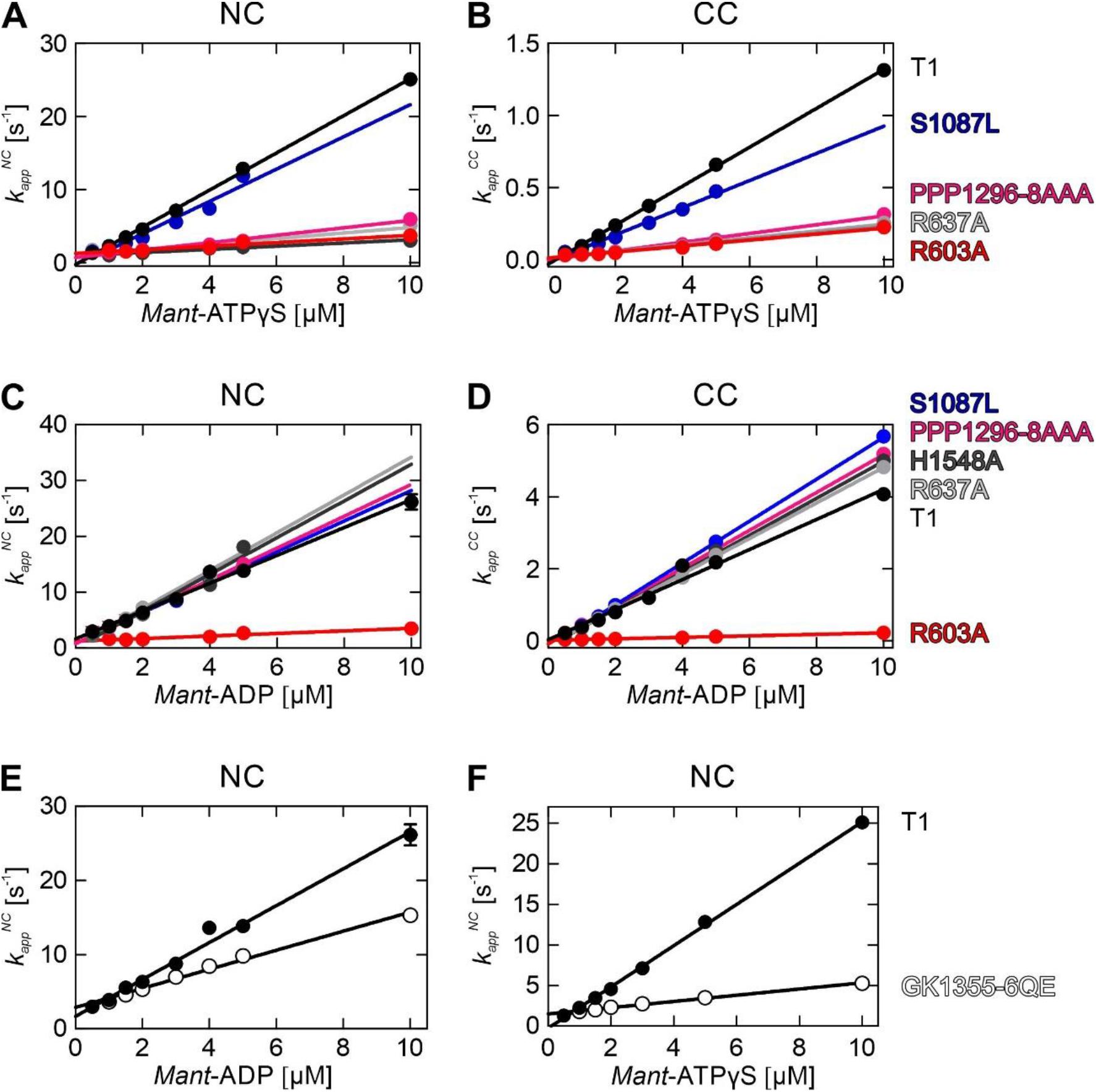
Effects of hBrr2^T1^ mutations on the kinetics of *mant*-ADP and *mant*-ATPγS binding. **A** and **B**. Apparent rate constants (*k_app_^NC^* and *k_app_^CC^*) of *mant*-ATPγS binding to the respective cassettes in hBrr2^T1^ and variants thereof. Binding of *mant*-ATPγS to either NC or CC is affected by all inter-cassette mutations tested. S1087L in the N-terminal HB domain originates from a retinitis pigmentosa-linked mutation of hBrr2, and was carried along as a negative control. **C** and **D**. Apparent rate constants (*k_app_^NC^* and *k_app_^CC^*) of *mant*-ADP binding to the respective cassettes of hBrr2^T1^ and variants thereof. While most inter-cassette mutants bind *mant*-ADP at similar rates as hBrr2^T1^, *mant*-ADP binding is almost completely abrogated at both NC and CC in R603A (red). hBrr2^T1^, black; S1087L (control with residue exchange in the N-terminal HB ratchet helix), blue; R603A (cassette interface NC), red; R637A (cassette interface NC), light gray; PPP1296-8AAA (linker), magenta; H1548A (cassette interface CC), dark gray. Coloring as in **A** and **B**. S1087L in the N-terminal HB domain originates from a retinitis pigmentosa-linked mutation of hBrr2, and was carried along as a negative control. **E** and **F**. Apparent rate constants (*k_app_^NC^*) of *mant*-ADP and *mant*-ATPγS binding to the NC of hBrr2^T1^ (closed circles) and its GK1355-6QE variant (altered CC nucleotide binding pocket; open circles). The time courses of nucleotide binding to GK1355-6QE were fitted by a single exponential indicating no detectable nucleotide binding at the CC, as expected due to the two residue exchanges in the CC binding pocket. As GK1355-6QE only has an intact NC nucleotide binding pocket, only the hBrr2^T1^ *k_app_^NC^* is shown for comparison. GK1355-6QE shows reduced rates for nucleotide binding at the NC, suggesting long range modulation of NC nucleotide binding by nucleotide binding at the CC in hBrr2^T1^. Values represent means ± SD of at least three independent measurements.

### Mutation of the C-terminal nucleotide binding pocket affects nucleotide binding at the N-terminal cassette

Mutations in motif I (Walker A motif or P-loop) of NTP-binding proteins and NTPases are known to interfere with nucleotide binding. In particular, a lysine residue in this motif is crucial for phosphate coordination and nucleotide stabilization (66). To test if nucleotide occupancy at the CC affects nucleotide binding at the NC, we generated a GK1355-6QE variant of hBrr2^T1^, bearing residue exchanges in motif I of the CC expected to abrogate nucleotide binding at the CC (Fig. 1C, right). Consistently, hBrr2^T1^ GK1355-6QE nucleotide association and dissociation time courses were best described by a single exponential, indicating nucleotide binding at the NC only. The association rate constants for *mant*-ADP and *mant*-ATPγS derived from the linear *k_app_^NC^* concentration dependence were more than 6-fold and about 2-fold slower, respectively, compared to the rates seen with wt hBrr2^T1^ (Fig. 6E,F; Table 1). The dissociation rate constants for ATPγS and ADP remained unchanged, as observed for the inter-cassette and linker mutants (Fig. 6E,F; Table 1). As a result, the NC nucleotide binding site of the GK1355-6QE mutant had about 2-fold and about 10-fold lower affinities for ADP and ATPγS, respectively, compared to the affinities displayed by the NC pocket of wt hBrr2^T1^. Thus, like the inter-cassette contacts, nucleotide binding at the CC also gears nucleotide affinities of the NC towards ATP.

### Molecular dynamics simulations suggest intra-molecular communication lines that might mediate long-range effects

Our rapid kinetics experiments indicated that the inter-cassette interface and linker between NC and CC influence nucleotide binding at both cassettes. Additionally, nucleotide binding to the CC can modulate the kinetics of ADP and ATP interaction with the NC, some 70 Å away. To delineate possible structural communication lines that could mediate these long-range effects, we conducted molecular dynamics (MD) simulations using available crystal structures of apo-hBrr2^T1^ (PDB ID 4F91) and hBrr2^T1^ with ADP bound at the NC and ATP bound at the CC (PDB ID 4F93) (50). By combining both crystal structures, we generated models of apo NC-CC (both nucleotide binding pockets empty), NC^ADP^-CC (ADP bound at the NC, CC empty), NC-CC^ATP^ (NC empty, ATP bound at the CC), NC^ADP^-CC^ATP^ (ADP bound at the NC, ATP bound at the CC) and NC^ATP^-CC^ATP^ (ATP bound at either cassette) and monitored their equilibrium dynamics at 300 K for 200 ns after equilibration.

Based on the MD trajectories, we first analyzed fluctuations in atomic positions. No significant differences in the fluctuations of Cα positions in the CC were observed when comparing trajectories of models bearing empty or nucleotide-bound CCs (Fig. 7A, right). In contrast, the region around residue 750 of the NC (RecA2 domain, region between motifs IV and IVa, involved in RNA binding) showed increased flexibility in NC^ADP^-CC and NC-CC^ATP^ (Fig. 7A, left). Fluctuations in this region were further increased when nucleotides were bound at both cassettes (NC^ADP^-CC^ATP^ and NC^ATP^-CC^ATP^; Fig. 7A, left). Detailed comparison of the linear correlation of fluctuations for the NC RecA1-RecA2-WH region confirmed an increased movement, manifested by stronger anti-correlations (darker red regions), between RecA2 and the other two domains (in particular the WH domain; Fig. 7B). The degree of this anti-correlation was increased in the fully-filled models (NC^ADP^-CC^ATP^ and NC^ATP^-CC^ATP^) compared to those with only one or no nucleotide bound (NC^ADP^-CC and NC-CC^ATP^; Fig. 7B). The anti-correlation was strongest for the NC^ATP^-CC^ATP^ model (Fig. 7B, boxed). Together, these observations indicate that nucleotide binding to the NC and CC leads to an additive increase in the flexibility of the region between motifs IV and IVa in the NC RecA2 domain.

**Fig. 7.**
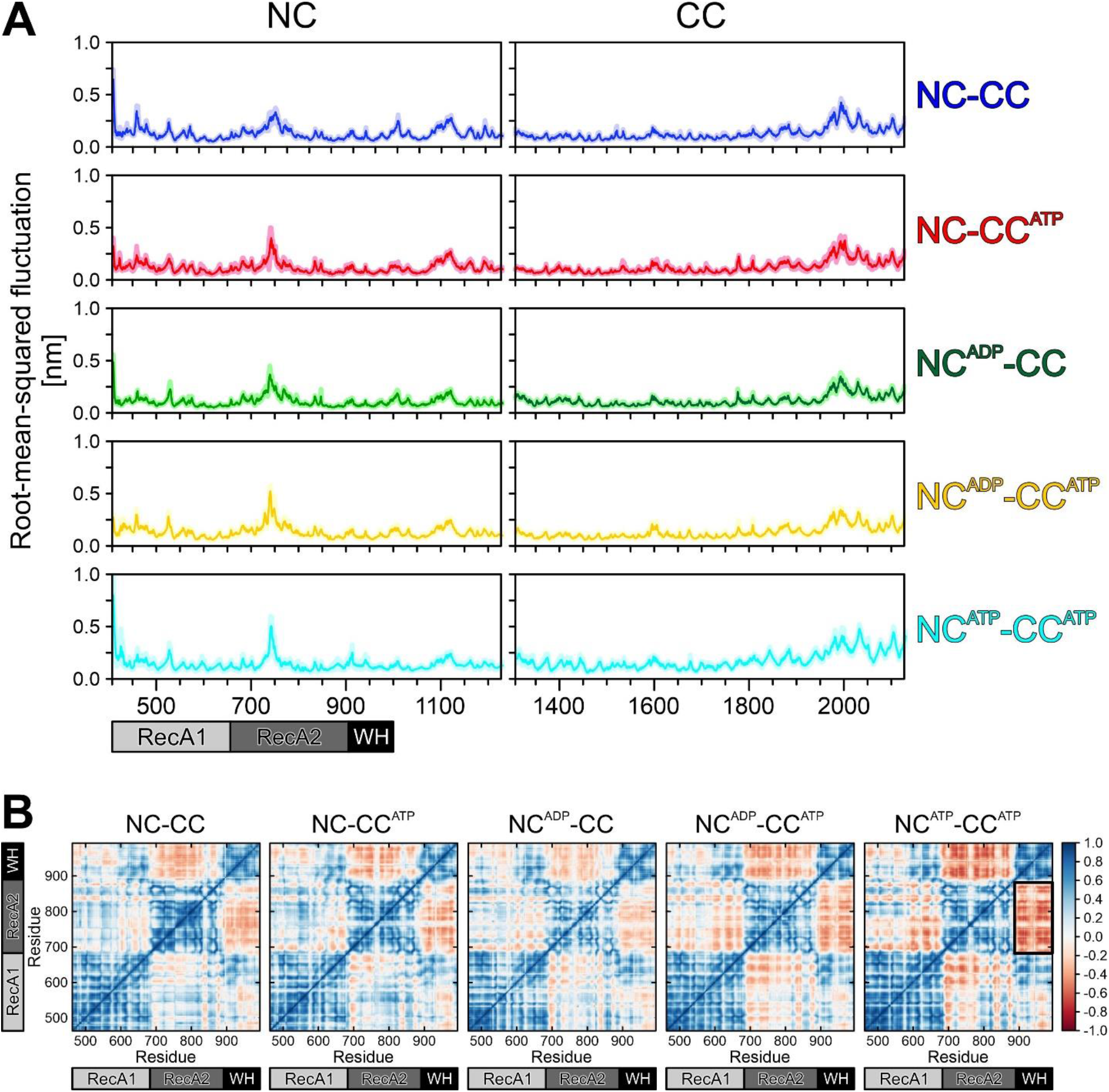
Brr2 flexibility. **A.** Flexibility of the NC (left) and of the CC (right) computed as root-mean-squared fluctuation. Solid lines, mean values; shaded areas, SEM, estimated from block averaging of the simulation data (see Methods). **B**. Linear correlation of positional fluctuations of Cα atoms of the RecA1, RecA2, and WH domains in the NC of hBrr2^T1^. Scale bar, degree of linear correlation.

We next quantified the distance distribution between G506 (motif I) and G854 (motif VI, involved in NTP binding/hydrolysis) in the NC as well as between corresponding residues in the CC (G1353 and G1689, respectively) as a measure for the widths of the nucleotide binding pockets in the MD trajectories of the various states. In the apo state, the NC pocket exhibited a bimodal width distribution with a predominant form around 7 Å and a minor form around 5 Å (Fig. 8A). Upon ADP binding to the NC alone (NC^ADP^-CC), only the 7 Å conformation remained (Fig. 8A). Upon additional binding of ATP to the CC (NC^ADP^-CC^ATP^), the NC pocket width was partially further increased, with a new distribution appearing at around 9 Å (Fig. 8A). With ATP bound at both cassettes (NC^ATP^-CC^ATP^), the NC nucleotide binding pocket showed a monomodal width distribution around 8 Å (Fig. 8A), *i.e.* intermediate between the widths seen in the bimodal NC^ADP^-CC^ATP^ distribution. ATP binding at the CC alone (NC-CC^ATP^) shifted the bimodal distribution in the NC of the apo state (5 and 7 Å) to larger widths (7 and 9 Å). As for the NC, the width of the CC nucleotide binding pocket was generally increased upon ATP binding (e.g. from around 4 Å in NC-CC to about 6 or 5 Å in NC-CC^ATP^ and NC^ATP^-CC^ATP^, respectively; Fig. 8B). ADP binding at the NC had particularly pronounced effects on the CC, increasing the nucleotide binding pocket of the latter cassette to about 8.5 Å in NC^ADP^-CC and to a bimodal 5/6 Å distribution in NC^ADP^-CC^ATP^ (Fig. 8B). These observations show that nucleotide-bound configurations of both cassettes are generally associated with larger widths of the respective nucleotide binding pockets compared to the unoccupied states. Moreover, ATP binding at the CC generally induces conformations of the NC associated with wider nucleotide-binding pockets. Likewise, ADP binding at the NC generally leads to a wider nucleotide binding pocket at the CC, while ATP at the NC might have a small effect in the opposite direction. Larger pockets and a trend towards bimodal distributions of the pocket widths upon nucleotide binding qualitatively agree with a higher flexibility of the motif IV-motif IVa region in the NC upon nucleotide binding (Fig. 7). Irrespective of the detailed effects, these results support the idea of structural communication between the two nucleotide binding pockets in hBrr2.

**Fig. 8.**
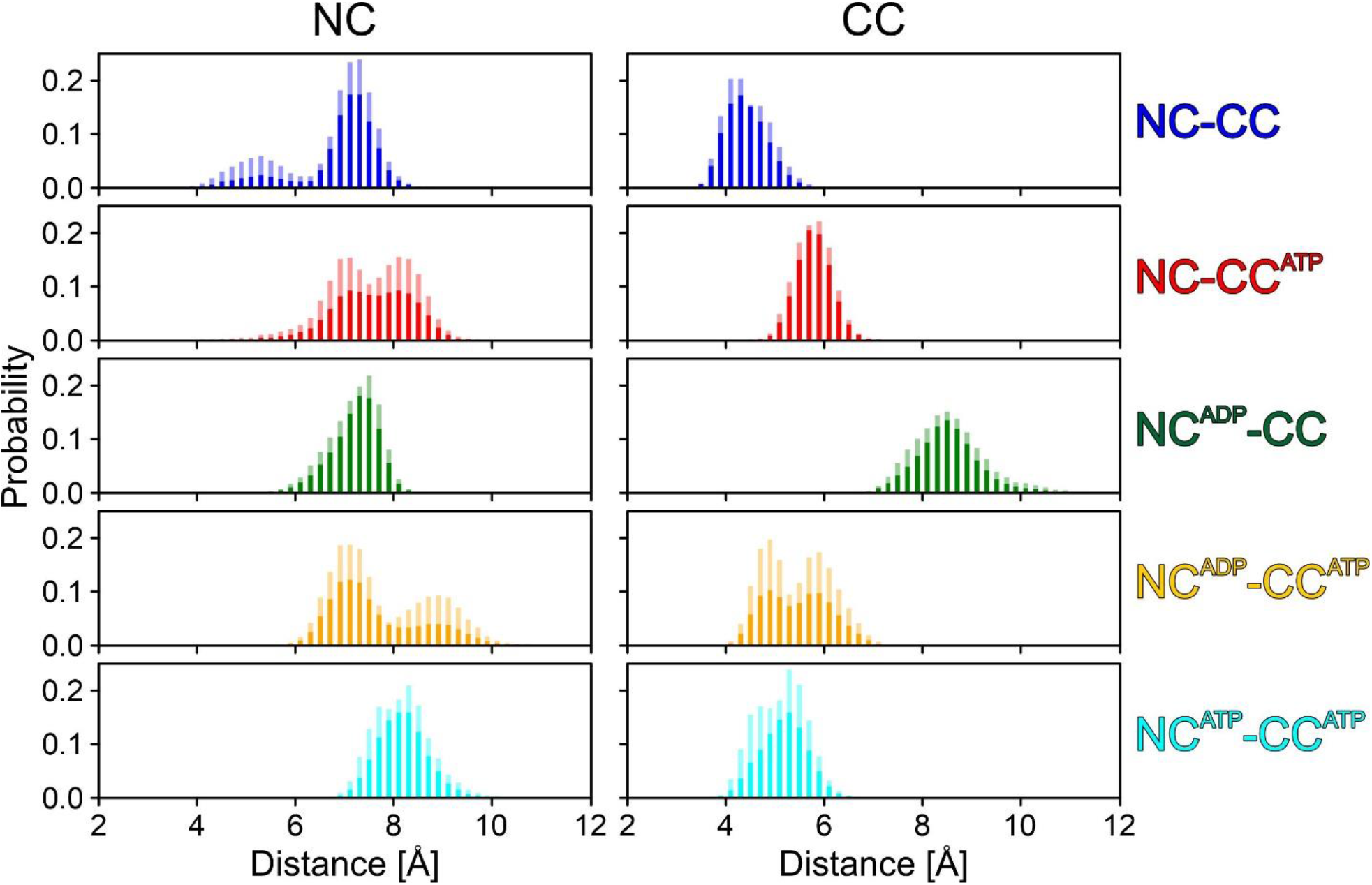
Widths of the nucleotide binding pockets. Widths of the nucleotide binding pockets of the NC, measured as the distance between G506 and G854 (left), and of the CC, measured as the distance between G1353 and G1689 (right). Solid bars, mean values; transparent bars, SEM, represented as mean + SEM.

To elucidate a possible mechanism of increased flexibilities upon NTP binding, hydrogen bond formation in the pockets with and without bound NTPs was analyzed. In the NC, the average number of hydrogen-bonded contacts between the protein and ADP or ATP was comparable (Table S2). Notably, residues binding the nucleotides did not engage in alternative, intra-hBrr2 hydrogen bonds in the absence of nucleotides. Thus, the observed increase in flexibility of the motif IV-motif IVa region in the NC upon nucleotide binding cannot be explained by redirected hydrogen bonding. However, nucleotide binding at the NC led to additional hydrogen bonds between K509 (motif I) and the backbone of A503 (ATP bound) or P504 (ADP bound), as well as between the side chain of N482 (Q-motif) and the backbone of A507 (motif I; ADP bound). In addition to cross-strutting by the bound nucleotide, such additional intra-hBrr2 hydrogen bonding might explain the increase in melting temperatures observed for all hBrr2 variants upon addition of ATP (Fig. S2). Most notably, N482 formed hydrogen bonds to Q485, when the CC was empty, *i.e.* to the glutamine of the Q-motif that binds the base of adenine nucleotides. Such sequestration of the Q-motif provides an explanation for our observation of reduced adenine nucleotide binding at the NC in the GK1355-6QE variant (Fig. 6E,F), in which nucleotide binding at the CC is abrogated.

Analysis of hydrogen bonds between the protein and the nucleotides in the CC showed an increased number of hydrogen bonds in NC^ADP^-CC^ATP^ and NC^ATP^-CC^ATP^ (7.1±1.1 and 6.8±1.1 hydrogen bonds, respectively) compared to the NC-CC^ATP^ state (4.0±0.9 hydrogen bonds; Table S2). Interestingly, G1353 (motif I), whose distance to G1689 (motif VI) we used as a measure of the pocket width, was involved in hydrogen bonds to ATP in NC^ADP^-CC^ATP^ and NC^ATP^-CC^ATP^. As in the NC, however, none of the residues binding the nucleotides *via* hydrogen bonds were engaged in intra-hBrr2 hydrogen bonds when the CC is unoccupied.

Finally, we sought to investigate the structural basis of long-range communication between the nucleotide binding pockets of the two cassettes, by which the CC pocket might influence nucleotide binding at the NC, and by which residues at the inter-cassette surfaces might exert their effects on nucleotide binding at either cassette. To this end, we extracted the shortest hydrogen-bonded paths, weighted by hydrogen-bond probabilities between residues along the path, between NC and CC nucleotide binding pockets from the MD trajectories of the various states (Fig. 9). Details of these shortest paths differed in the different MD trajectories. Model NC^ATP^-CC^ATP^ exhibited the lowest diversity in shortest paths between the pocket residues (Fig. 9A). The helix containing T1578 (following motif IV^CC^) was only involved in shortest paths in models NC-CC^ATP^ and NC^ADP^-CC^ATP^, whereas the helix that contains H1548 (preceding motif IV^CC^) was part of shortest paths connecting the pocket residues in all models except for NC^ADP^-CC^ATP^ (Table 3). In the two models with ADP bound at the NC, *i.e.* NC^ADP^-CC and NC^ADP^-CC^ATP^, the shortest paths from K509 and T510 (motif I^NC^) that ran along the backbone of the helix following motif I^NC^ through a strong hydrogen bond between R647 (preceding motif III^NC^) and E497 (between Q motif^NC^ and motif I^NC^) would provide a short-cut to motif III^NC^. The hydrogen bond-based communication between the two cassettes appears to be dominated by the highly probable hydrogen bonds between R603 (between motifs Ic^NC^ and II^NC^) and D1575 (helix following motif IV^CC^), between R637 (between motifs II^NC^ and III^NC^) and D1583 (helix following motif IV^CC^) and between E602 (between motifs Ic^NC^ and II^NC^) and K1544 (preceding motif IV^CC^; Table S3). In all models, except for NC^ATP^-CC^ATP^, at least one shortest path between the nucleotide binding sites runs *via* R637 (Table 3). It is interesting to note that R637-D1583 is the interface crossing in some paths of the models with empty CC (Table S4), although in these models this hydrogen bond is less probable than in the other models (Table S3). In the NC-CC^ATP^ model the contrary is observed: the interface is crossed between R637 and D1583 (Table S4), which form a hydrogen bond throughout the course of the simulation, but residue 1583 is not contained in the shortest paths between active sites (Table 3). The interface is crossed from E602 to K1544 (Table S4), which is also the crossing point for model NC^ATP^-CC^ATP^, although this hydrogen bond has only low probability in this model (Table S3). R603 is part of the shortest paths between nucleotide pockets in the NC^ADP^-CC^ATP^ and NC-CC models (Table 3). In the NC^ADP^-CC^ATP^ model, R603 is part of the shortest paths between almost each pair of active site residues analyzed (Table 3). H1548 is involved in all shortest paths in the NC-CC^ATP^ model and in some shortest paths in the NC^ATP^-CC^ATP^ model (Table 3). These paths cross the interface between E602 and K1544 (Table S4). It is interesting to note that models with ATP bound to the CC show shortest paths crossing the interface at a similar location: models NC-CC^ATP^ and NC^ADP^-CC^ATP^ at E602 (to K1544 and subsequently to H1548) and model NC^ADP^-CC^ATP^ at the neighboring R603 (to L1540 or D1575; Table S4).

**Fig. 9.**
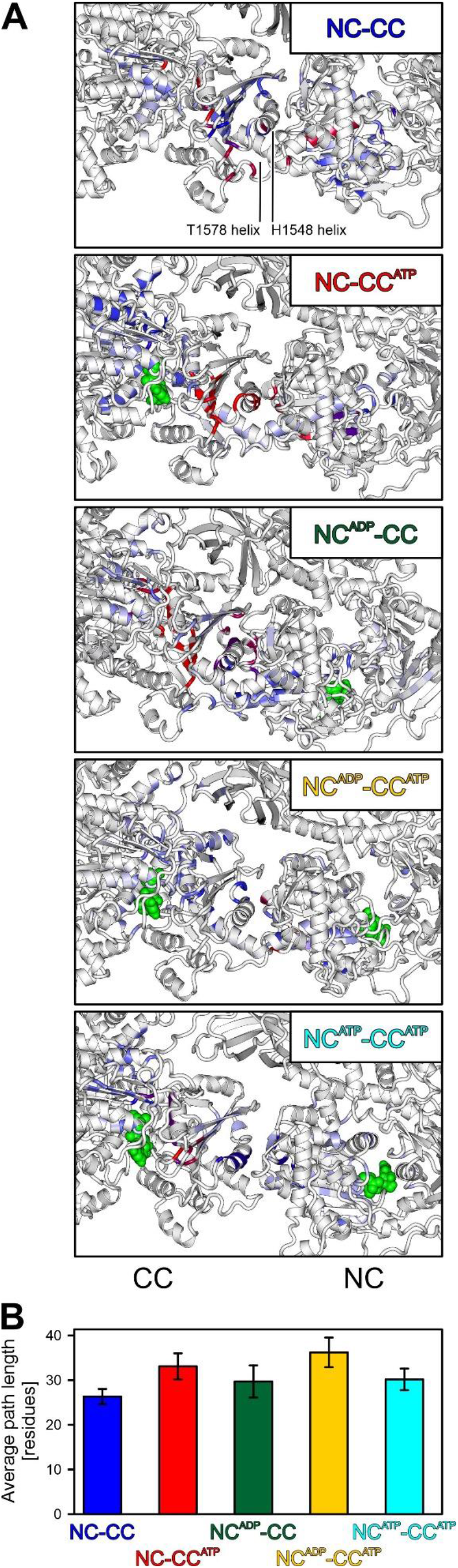
Shortest paths. **A.** Shortest communication paths visualized by highlighting the residues participating in the *k*-shortest pathways between the nucleotide binding pockets of NC and CC by frequency (low-to-high, blue-to-red). Green spheres, bound nucleotides. **B.** Average path lengths (number of connected residues) of shortest communication paths between the nucleotide binding pockets of NC and CC. Values represent means ± SEM.

**Table 3.** Participation of mutated residues and of E602 and H1534 in the *k*-shortest paths (*k*=10).

| Path | NC-CC | NC-CC<sup>ATP</sup> | NC<sup>ADP</sup>-CC | NC<sup>ADP</sup>-CC<sup>ATP</sup> | NC<sup>ATP</sup>-CC<sup>ATP</sup> |
| --- | --- | --- | --- | --- | --- |
| **E616-E1455** | R637 | E602, R637,<br>K1544, H1548 | E602, R637,<br>H1534 | R603 | E602, K1544,<br>H1548 |
| **K509-K1356** | R637 | E602, K1544,<br>H1548, T1578 | E602, H1534 | E602, R603 |  |
| **N820-N1692** | R603, H1534 | E602, K1544,<br>H1548 | E602, R637,<br>H1534 | R603 | E602, K1544,<br>H1548 |
| **Q484-Q1332** | E602, R637,<br>H1534 | E602, K1544,<br>H1548 | E602, H1534,<br>F1717 | R603 | E602, K1544,<br>H1548 |
| **R855-E1455** | R603, R637 | E602, K1544,<br>H1548 | E602, R637,<br>H1534 | R603, R637 | E578, H1534 |
| **T510-T1357** | R637 | K1544,<br>H1548, T1578 | E602, R1195,<br>H1534 | R603 | E602, K1544,<br>H1548 |

Together, these observations show that depending on the nucleotide occupation of the cassettes their inter-cassette contacts change, providing a molecular basis for the observation that mutations of residues at the cassette interface impact binding of ATPγS to either cassette. The additional reduced ADP binding to both cassettes upon R603A mutation is furthermore reflected in the importance of R603 in shortest paths of the ADP-bound model NC^ADP^-CC^ATP^ (Table 3).

## DISCUSSION

Results presented here reveal intricate networks of intra-molecular communication in the Brr2 RNA helicase that modulate the adenine nucleotide binding specificities and affinities of the enzyme’s N- and C-terminal nucleotide binding pockets. Changes at inter-cassette contacts or in a linker connecting the two cassettes affect nucleotide binding at both pockets. Furthermore, the N-terminal nucleotide binding pocket can “sense” nucleotide binding at the C-terminal pocket, *i.e.* over a distance of about 70 Å. Brr2 architecture and dynamics suggest likely molecular communication lines that mediate these long-range effects, which run through layers of structural elements in the vicinity of the nucleotide binding pockets, and which connect the two nucleotide binding pockets across the inter-cassette interface.

### Mechanism of nucleotide binding by Brr2

To date, there are a limited number of reports on kinetic measurements of nucleotide binding to RNA and DNA helicases. To our knowledge, our work represents the first transient kinetics analysis of nucleotide binding to a double-cassette Ski2-like helicase. Due to the presence of two nucleotide binding pockets in Brr2, we initially characterized the kinetics of *mant*-ADP and *mant*-ATPγS binding and dissociation using single cassette constructs, hBrr2^NC^ and hBrr2^CC^. Detailed characterization of the isolated cassettes allowed us to subsequently unequivocally assign double exponentials observed upon nucleotide binding or dissociation to/from dual-cassette constructs of hBrr2 to either of the cassettes.

*Mant*-nucleotide binding to hBrr2 generates FRET from NC and CC tryptophans to the *mant* moiety of the bound nucleotide. hBrr2 contains one and two trytophans within about 20 Å distance of nucleotides bound at the NC and CC, respectively (Fig. S1). The differences in the magnitudes of the FRET signals seen with *mant*-ADP or *mant*-ATPγS, with *mant*-ADP binding resulting in a higher amplitude compared to *mant*-ATPγS binding (Fig. 2A,B), are explained by lower occupancy of the nucleotide binding pockets in the case of *mant*-ATPγS, due to a lower affinity of ATPγS compared to ADP (Table 1). Higher affinity for ADP is also reflected in the better defined electron densities for ADP/*mant*-ADP compared to *ATPγS/mant*-ATPγS in the corresponding crystal structures (Fig. 3).

Nucleotide binding to other helicases such as Rep, DnaB and RecG showed multi-phasic kinetics, minimally characterized by a two-step mechanism, with a rapid nucleotide binding phase followed by nucleotide accommodation that resulted in a state of high nucleotide affinity characterized by high fluorescence (67-69). For hBrr2, both in the single-cassette as well as in the double-cassette constructs, nucleotide binding at both pockets could be modeled as a single-step, reversible process, as also observed for the DbpA helicase (70).

### Implications for the differential functions of the two helicase cassettes in Brr2

Our results demonstrate that, in the absence of RNA, *mant*-ATPγS binds more weakly to both cassettes than *mant*-ADP as also observed for the DEAD-box RNA helicase, DbpA (70). In addition, a comparison of the kinetics of nucleotide binding by the hBrr2 cassettes indicates that hBrr2^NC^ binds and releases nucleotides faster than hBrr2^CC^, both in isolation as well as in the dual-cassette constructs tested (Fig. 4 and 5). The high hBrr2^CC^ nucleotide affinity is the result of a very low nucleotide dissociation rate, with *mant*-ADP and *mant*-ATPγS dissociation being 945 and 800 times slower for hBrr2^CC^ compared to hBrr2^NC^, respectively.

These results are consistent with the functions of the two cassettes for the activity of the enzyme. Fast nucleotide dynamics at the NC are in line with the NC being the active helicase cassette in the protein, which undergoes rounds of nucleotide binding, hydrolysis and release of the products, and which couples these transactions to conformational changes that give rise to translocation of the enzyme on the substrate RNA. The CC, in contrast, is inactive as an enzyme but stimulates the helicase activity of the NC (50). Rather than relying on conformational changes driven by nucleotide transactions, the CC may remain permanently bound to a nucleotide during Brr2-mediated RNA unwinding and provide a stable scaffold, which offers anchoring points for the NC, and can thereby support productive cycles of conformational changes in the NC. This picture is fully in line with our finding that mutations in inter-cassette contacts and in the inter-cassette linker, which would in part abrogate the ability of the NC to take advantage of the CC scaffold to transition between conformational states, influence nucleotide binding at the NC (Fig. 6). Furthermore, mutations in the CC nucleotide binding pocket designed to abrogate nucleotide binding also resulted in reduced nucleotide binding rates at the NC (Fig. 6).

Previously, mutations in the inter-cassette interface were seen to have only mild effects on the RNA-stimulated ATPase activity of hBrr2 (50). These observations could indicate that, in the presence of RNA, the impact of the mutations on nucleotide binding is not as pronounced as observed in our setup. However, some of the mutations still gave rise to significant defects in the RNA unwinding activity of the respective hBrr2 variants (50). Therefore, we expect that the mutations interfere with the coupling of the ATPase to the helicase activity in the NC.

## EXPERIMENTAL PROCEDURES

### Cloning and mutagenesis

Codon-optimized DNA fragments encoding hBrr2^FL^ (residues 1-2136) and fragments thereof (hBrr2^T1^: residues 395-2129; hBrr2^NC^: residues 395-1324; hBrr2^CC^: residues 1282-2136) were cloned into a modified pFL vector (EMBL, Grenoble) to produce proteins with a TEV-cleavable N-terminal His_10_-tag (50). Site-directed mutagenesis was performed using the QuikChange II XL Site-Directed Mutagenesis Kit (Stratagene). All constructs were verified by sequencing. All plasmids were transformed into *Escherichia coli* DH10MultiBacY cells (provided by Imre Berger, University of Bristol) and further integrated *via* Tn7 transposition into the baculovirus genome (EMBacY) maintained as a bacterial artificial chromosome (BAC) (71). The Tn7 transposition site was embedded in a *lacZα* gene allowing the selection of positive EMBacY recombinants *via* blue/white screening. Recombinant BACs were isolated from the bacterial hosts and used to transfect Sf9 cells (Invitrogen).

### Protein production

All proteins were produced by recombinant baculoviruses in insect cells, as described previously (50,61). Briefly, for initial virus (V_0_) production, the isolated recombinant EMBacY was transfected into adhesive Sf9 cells (Invitrogen) in 6-well plates. The efficiency of transfection was monitored by eYFP fluorescence. The V_0_ virus generation was used to infect 50 ml Sf9 cells for virus amplification. The second, high titer virus generation (V_1_) was then used to infect 1200 ml High Five™ cells (Invitrogen) for large scale protein production. The infected cells were harvested when the eYFP signal reached a plateau and before the cell viability dropped below 90 %.

### Protein purification

Proteins were purified as described previously (50,61). Briefly, for protein production for biochemical/biophysical experiments, the High Five™ cell pellet was resuspended in 50 mM HEPES-NaOH, pH 8.0, 600 mM NaCl, 2 mM β-mercaptoethanol, 0.05 % NP40, 1.5 mM MgCl_2_, 20 (v/v) % glycerol, 10 mM imidazole, supplemented with EDTA-free protease inhibitor (Roche) and lyzed by sonication using a Sonopuls Ultrasonic Homogenizer HD 3100 (Bandelin). The target was captured from the cleared lysate on a 5 ml HisTrap FF column (GE Healthcare) and eluted with a linear gradient from 10 to 250 mM imidazole. The eluted protein containing the protein of interest was diluted to a final concentration of 80 mM NaCl, treated with RNaseA (Sigma) and loaded on a Mono Q 10/100 GL column (GE Healthcare) equilibrated with 50 mM Tris-HCl, pH 8.0, 50 mM NaCl, 5 (v/v) % glycerol, 2 mM β-mercaptoethanol. The protein was eluted with a linear 0.05 to 1.5 M NaCl gradient and further purified by gel filtration on a HiLoad Superdex 200 16/60 column (GE Healthcare) in 40 mM Tris-HCl, pH 8.0, 200 mM NaCl, 20 (v/v) % glycerol, 2 mM DTT. The peak fractions were concentrated, flash-frozen in liquid nitrogen and stored at −80 °C.

For protein production for crystallization, the hBrr2^T1^ insect cell pellet was lysed by sonification for 30 min in 50mM HEPES-NaOH pH 7.5, 600mM NaCl, 10% (w/v) glycerol, 0.05% (v/v) Nonidet P-40, 20μg/ml DNase I, 2mM β-mercaptoethanol, containing Complete EDTA-free protease inhibitors. After centrifugation and loading onto a HisTrap FF column, the protein was eluted with 250 mM imidazole. TEV protease was added for cleavage of the His-tag, and the mixture was dialyzed overnight in 40 mM HEPES-NaOH, pH 7.5, 500 mM NaCl, 10 % (w/v) glycerol, 15 mM imidazole, 2 mM β-mercaptoethanol. The cleaved protein was collected in the flow-through of a HisTrap FF column. After five-fold dilution with 25 mM Tris-HCl, pH 8.0, 50 mM NaCl, 5 % (v/v) glycerol, 2 mM DTT and treatment with RNase A, the protein was loaded on a HiPrep Heparin FF column (GE Healthcare) equilibrated with 25 mM Tris-HCl, pH 8.0, 50 mM NaCl, 5 % (v/v) glycerol, 2 mM DTT, and eluted by a linear increase of NaCl to 750 mM. The fractions of interest were combined and chromatographed on a HiLoad Superdex 200 16/60 column in 10 mM Tris-HCl, pH 7.5, 200 mM NaCl, concentrated to 10 mg/ml, flash-flozen in liquid nitrogen and stored at −80 °C.

The hJab1 ^ΔC^ insect cell pellet was lysed by sonification for 30 min in 50 mM Tris-HCl, pH 8.0, 300 mM NaCl, 5 % (v/v) glycerol, 0.05 % (v/v) NP-40, 2 mM DTT, supplemented with Complete EDTA-free protease inhibitors. After centrifugation, the protein was captured on glutathione sepharose beads (GE Healthcare) and eluted with 10 mM reduced glutathione. Buffer was exchanged to 50 mM Tris-HCl, pH 8.0, 300 mM NaCl, 5 % (v/v) glycerol, 2 mM DTT on a HiLoad Superdex 75 26/60 column (GE Healthcare). After treatment with Prescission protease overnight, the hJab1^ΔC^ protein lacking the GST tag was collected in the flow-through of glutathione sepharose beads. Subsequently, the protein was loaded on a HiLoad Superdex 75 16/60 column in 10 mM Tris-HCl, pH 8.0, 150 mM NaCl, concentrated to 4 mg/ml, flash-flozen in liquid nitrogen and stored at −80 °C.

For complex formation, hBrr2^T1^ was combined with a 1.5-fold molar excess of hJab1^ΔC^, and the complex was purified by gel filtration on a Superdex 200 10/300 global increase column (GE Healthcare) in 20 mM Tris-HCl, pH 8.0, 150 mM NaCl, concentrated to 6 mg/ml, flash-frozen in liquid nitrogen and stored at −80°C.

### Crystallographic analyses

Crystals of the hBrr2^T1^-hJab1 ^ΔC^ complex were grown in a 24-well plate with a reservoir solution of 0.1 M HEPES-NaOH, pH 8.0, 0.1 M MgCl_2_, 8 % PEG 3350, as described before (61). Crystals were soaked for 1 h in 10 mM ADP or ATPγS, or in 1 mM *mant*-ADP or *mant*-ATPγS (Jena Bioscience) in reservoir solution. After cryoprotection with 25 % (v/v) ethylene glycol in reservoir solution, crystals were flash-cooled in liquid nitrogen.

Diffraction data were acquired at beamline 14.2 of the BESSY II storage ring (Berlin, Germany) and processed with XDS (72). Molecular replacement was done with Phenix (73) using the structure coordinates of the hBrr2^T1^-hJab1 ^ΔC^ complex (PDB ID 6S8Q) (61). The structures were manually adjusted with Coot (74) and automatically refined with Phenix. Restraints for the ligands were generated by the “eLBOW”-tool of Phenix. Structure figures were prepared with PyMOL (Version 1.8 Schrödinger, LLC).

### Characterization of protein variants

DSF experiments were done in a 96-well plate in a plate reader combined with a thermocycler (Stratagene Mx3005P). All hBrr2 constructs were diluted to a final concentration of 3 μM in purification buffer (± 2 mM ATP/Mg^+2^) supplemented with 10×SYPRO orange (1:500 dilution of the stock) in a total volume of 20 μl. The temperature was increased linearly from 25 °C to 95 °C and the fluorescence emission was monitored in steps of 1 °C/min with hold steps of 30 s between reads. The fluorescence intensity was then plotted as a function of temperature. The sigmoidal curve from each construct was normalized and corrected for the background signal of the fluorophore in the buffer. The inflection points of the curves, representing the thermal melting temperature, were extracted from the first derivatives of the melting profiles using OriginLab.

### Rapid kinetic measurements

The kinetics of the interaction of hBrr2 variants with nucleotides were characterized *via* fluorescence stopped-flow measurements on an SX-20MV spectrometer (Applied Photophysics). The fluorescence of *mant*-labeled nucleotides was excited using 290 nm light *via* FRET from tryptophan residues in the proximity of the nucleotide binding pockets and measured at 90 ° after passing a cut-off filter (KV 408, Schott). FRET was observed only when both donor and acceptor were present since a negligible fluorescence change of tryptophan was observed when non-fluorescent nucleotides were bound to the proteins. Association experiments were performed by rapidly mixing equal volumes (60 μl) of the reactants (syringe 1 contained hBrr2 variants while syringe 2 contained the *mant* nucleotides) in 20 mM HEPES-NaOH, pH 8.0, 150 mM NaCl, 1.5 mM MgCl_2_ at 20 °C and monitoring fluorescence change over time. Dissociation or chase experiments were performed similarly by rapidly mixing equal volumes (60 μl) of the reactants (syringe 1 contained hBrr2 variants in complex with *mant* nucleotides while syringe 2 contained an excess of unlabeled nucleotides) at 20 °C and monitoring fluorescence change over time. In all cases, 1000 data points were acquired in logarithmic sampling mode. The data were visualized using the Pro-Data Viewer software package (Applied Photophysics). The final curves were obtained by averaging 7-10 individual traces after normalizing each data point to initial F_0_. Data were evaluated by fitting to a single exponential function with a characteristic time constant (*k_app_*), amplitude (*F_1_*) and another variable for the final signal (*F_∞_*) according to the equation, *F* = *F_∞_* + *F_1_exp(-k_app_t*), in which *F* is the fluorescence at time *t*. For constructs containing two nucleotide binding sites, two exponential terms were used with two characteristic time constants (*k_app1_*, *k_app2_*), amplitudes of the signal change (*F_1_*, *F_2_*) and another variable for the final signal (*F_∞_*) according to the equation, *F* = *F_∞_* + *F_1_*exp(-*k_app1_*t) + *F_2_*exp(-*k_app2_*t). Dependencies of the apparent rate constants on nucleotide concentration were fitted by a linear equation, *k_app_* = *k_1_*[mant-nucleotide]+*k-_1_*, in which *k_1_* represents the nucleotide association rate constant (derived from the slope), and *k-_1_* represents the nucleotide dissociation rate constant (derived from the Y-axis intercept). Calculations and statistical analysis were performed using Prism software (GraphPad).

### Molecular dynamics simulations

hBrr2^403 to 2125^ was modeled based on PDB entries 4F91 (for the apo form) and 4F93 (for nucleotide-bound forms) (50). ADP was changed *in silico* to ATP and one magnesium ion was modeled into the NC binding pocket using its position relative to ATP in the CC pocket as a template. Protein, nucleotides and ions were described with the CHARMM force field (75). The (nucleotide-bound) proteins were first relaxed by 500 steps of steepest descend. The relaxed structures were then solvated by TIP3P water (76) in a dodecahedral box, extending 1.5 nm from the solute (~ 1.8 nm length). After 5000 steps of optimization, keeping the nucleotide fixed, 1 ns of molecular dynamics simulations with all protein heavy atoms positionally restrained were performed to further equilibrate the systems at 300 K, using a Berendsen thermostat (77). Finally, molecular dynamics production runs of 200 ns length were performed in an NVT ensemble, at 300 K controlling temperature by canonical sampling through canonical velocity-rescaling (78), and 2 fs time steps for the integration with the LINCS algorithm (79) to constrain covalent bonds. Electrostatic interactions were treated with the particle mesh Ewald method (80) on a grid with 0.16 spacing and a short-range cut-off of 1.4 nm. The same cut-off was applied to the van der Waals interactions. All simulations were performed with Gromacs 4.6.7 (81).

For the analyses, only the last 100 ns simulation time were considered. Hydrogen bonds were defined geometrically by a 3.2 Å maximal distance between donor (D) and acceptor (A) atoms and 42° maximum deviation from linearity for the D-H···A angle.

For the communication analysis, a weighted graph was constructed, in which the protein residues form the nodes, and edges between nodes were defined by the occurrence of a hydrogen bond in the course of the simulation. The probabilities of these hydrogen bonds to occur served as edge weights. The probability of hydrogen bonds between two residues i and j, HB_ij_, were converted into communication costs C_ij_ = -ln(HB_ij_). Consecutive residues, *i.e.* covalently bound, were assigned a communication cost of zero. Shortest paths were selected from the resulting communication graph using Dijkstra’s algorithm (82), with the path lengths taken as the sum of the edge weights along the path. The analysis of hydrogen bonds, setup and evaluation of communication graphs were carried out using our own Java code, based on the Jgraph library (83).

Protein flexibilities were analyzed using the rmsf tool of the Gromacs program. Errors were estimated from block averaging, partitioning the last 100 ns of the simulation data into to five blocks of 20ns length each.

## Acknowledgments

We thank Imre Berger, University of Bristol, for *E. coli* DH10MultiBacY cells. We acknowledge access to beamlines BL14.1/2/3 of the BESSY II storage ring (Berlin, Germany) *via* the Joint Berlin MX-Laboratory sponsored by the Helmholtz-Zentrum Berlin für Materialien und Energie, Freie Universität Berlin, Humboldt-Universität zu Berlin, the Max-Delbrück Centrum für Molekular Medizin, the Leibniz-Forschungsinstitut für Molekulare Pharmakologie and Charité – Universitätsmedizin Berlin. We are grateful for Computational resources provided by the North-German Supercomputing Alliance (HLRN). This work was funded by grant TRR186/A15 from the Deutsche Forschungsgemeinschaft to MCW. KFS was supported by a Dahlem International Network PostDoc Fellowship from Freie Universität Berlin.

## Footnotes

This work was funded by grant TRR186/A15 from the Deutsche Forschungsgemeinschaft to MCW. KFS was supported by a Dahlem International Network PostDoc Fellowship from Freie Universität Berlin.

The abbreviations used are: BAC, bacterial artificial chromosome; CC, C-terminal cassette; DSF, differential scanning fluorimetry; DTT, dithiothreitol; FRET, fluorescence resonance energy transfer; h, human; HB, helical bundle domain; HLH, helix-loop-helix domain; IG, immunoglobulin-like domain; *mant*, methylanthraniloyl; MD, molecular dynamics; NC, N-terminal cassette; NTR, N-terminal region; NTPase, nucleic acid-dependent nucleotide tri-phosphatase; pre-mRNA, precursor messenger RNA; RNP, ribonucleoprotein complex; SEM, standard error of the mean; SF2, superfamily 2; sn, small nuclear; ss, single-stranded; T1, truncation 1; Tris, tris(hydroxymethyl)aminomethane; v/v, volume/volume; WH, winged-helix domain; wt, wild type.

## SUPPLEMENTAL MATERIAL

### SUPPLEMENTAL TABLES

**Table S1.**
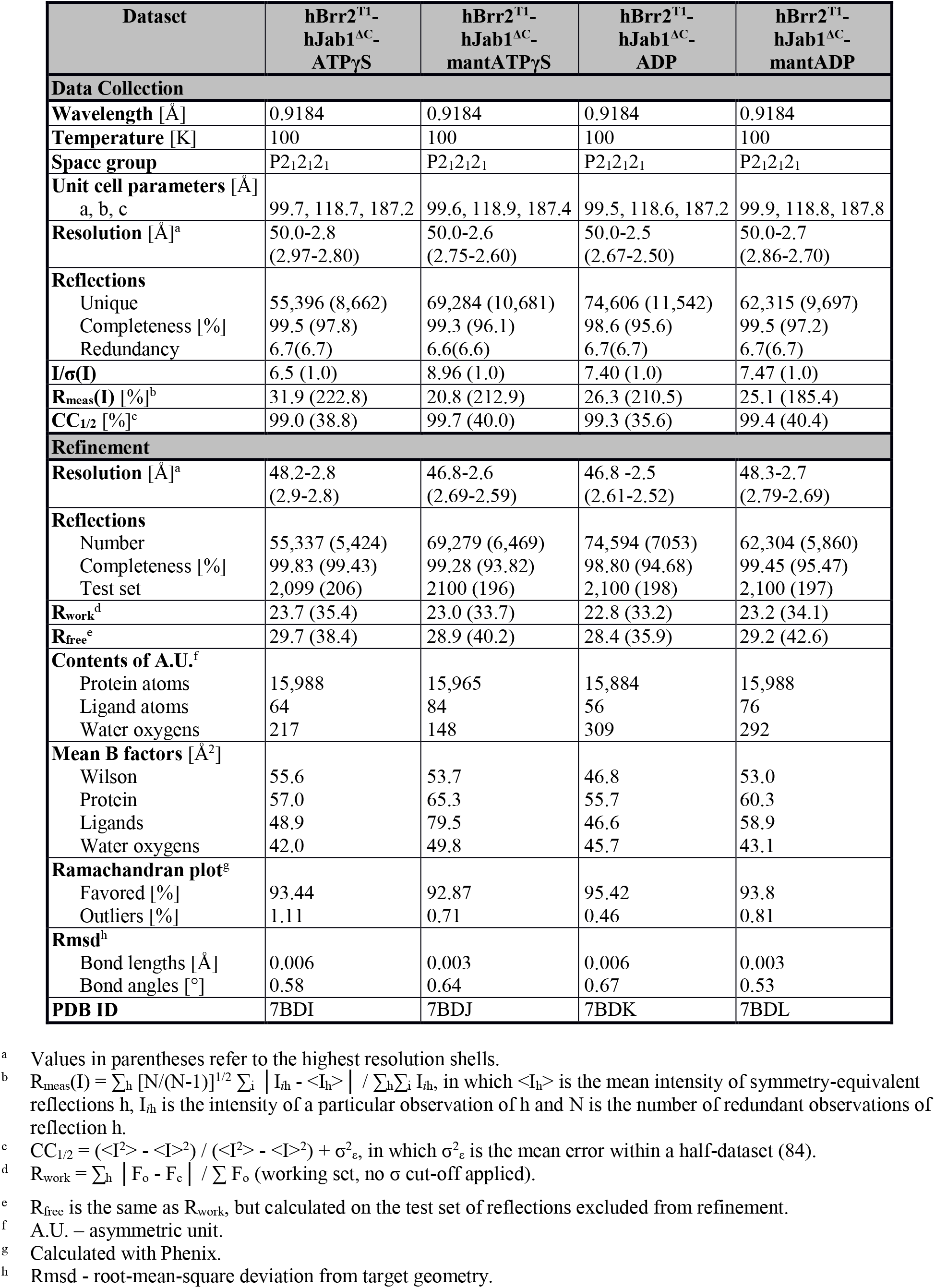
Crystallographic data.

| Dataset | hBrr2<sup>T1</sup>-<br>hJab1<sup>ΔC</sup>-<br>ATP<sub>γ</sub>S | hBrr2<sup>T1</sup>-<br>hJab1<sup>ΔC</sup>-<br>mantATP<sub>γ</sub>S | hBrr2<sup>T1</sup>-<br>hJab1<sup>ΔC</sup>-<br>ADP | hBrr2<sup>T1</sup>-<br>hJab1<sup>ΔC</sup>-<br>mantADP |
| --- | --- | --- | --- | --- |
| **Data Collection** |  |  |  |  |
| **Wavelength** [Å] | 0.9184 | 0.9184 | 0.9184 | 0.9184 |
| **Temperature** [K] | 100 | 100 | 100 | 100 |
| **Space group** | P2<sub>1</sub>2<sub>1</sub>2<sub>1</sub> | P2<sub>1</sub>2<sub>1</sub>2<sub>1</sub> | P2<sub>1</sub>2<sub>1</sub>2<sub>1</sub> | P2<sub>1</sub>2<sub>1</sub>2<sub>1</sub> |
| **Unit cell parameters** [Å]<br>a, b, c | 99.7, 118.7, 187.2 | 99.6, 118.9, 187.4 | 99.5, 118.6, 187.2 | 99.9, 118.8, 187.8 |
| **Resolution** [Å]<sup>a</sup> | 50.0-2.8<br>(2.97-2.80) | 50.0-2.6<br>(2.75-2.60) | 50.0-2.5<br>(2.67-2.50) | 50.0-2.7<br>(2.86-2.70) |
| **Reflections**<br>Unique<br>Completeness [%]<br>Redundancy | 55,396 (8,662)<br>99.5 (97.8)<br>6.7(6.7) | 69,284 (10,681)<br>99.3 (96.1)<br>6.6(6.6) | 74,606 (11,542)<br>98.6 (95.6)<br>6.7(6.7) | 62,315 (9,697)<br>99.5 (97.2)<br>6.7(6.7) |
| **I/σ(I)** | 6.5 (1.0) | 8.96 (1.0) | 7.40 (1.0) | 7.47 (1.0) |
| **R<sub>meas</sub>(I)** [%]<sup>b</sup> | 31.9 (222.8) | 20.8 (212.9) | 26.3 (210.5) | 25.1 (185.4) |
| **CC<sub>1/2</sub>** [%]<sup>c</sup> | 99.0 (38.8) | 99.7 (40.0) | 99.3 (35.6) | 99.4 (40.4) |
| **Refinement** |  |  |  |  |
| **Resolution** [Å]<sup>a</sup> | 48.2-2.8<br>(2.9-2.8) | 46.8-2.6<br>(2.69-2.59) | 46.8-2.5<br>(2.61-2.52) | 48.3-2.7<br>(2.79-2.69) |
| **Reflections**<br>Number<br>Completeness [%]<br>Test set | 55,337 (5,424)<br>99.83 (99.43)<br>2,099 (206) | 69,279 (6,469)<br>99.28 (93.82)<br>2100 (196) | 74,594 (7053)<br>98.80 (94.68)<br>2,100 (198) | 62,304 (5,860)<br>99.45 (95.47)<br>2,100 (197) |
| **R<sub>work</sub>**<sup>d</sup> | 23.7 (35.4) | 23.0 (33.7) | 22.8 (33.2) | 23.2 (34.1) |
| **R<sub>free</sub>**<sup>e</sup> | 29.7 (38.4) | 28.9 (40.2) | 28.4 (35.9) | 29.2 (42.6) |
| **Contents of A.U.**<sup>f</sup><br>Protein atoms<br>Ligand atoms<br>Water oxygens | 15,988<br>64<br>217 | 15,965<br>84<br>148 | 15,884<br>56<br>309 | 15,988<br>76<br>292 |
| **Mean B factors** [Å<sup>2</sup>]<br>Wilson<br>Protein<br>Ligands<br>Water oxygens | 55.6<br>57.0<br>48.9<br>42.0 | 53.7<br>65.3<br>79.5<br>49.8 | 46.8<br>55.7<br>46.6<br>45.7 | 53.0<br>60.3<br>58.9<br>43.1 |
| **Ramachandran plot**<sup>g</sup><br>Favored [%]<br>Outliers [%] | 93.44<br>1.11 | 92.87<br>0.71 | 95.42<br>0.46 | 93.8<br>0.81 |
| **Rmsd**<sup>h</sup><br>Bond lengths [Å]<br>Bond angles [°] | 0.006<br>0.58 | 0.003<br>0.64 | 0.006<br>0.67 | 0.003<br>0.53 |
| **PDB ID** | 7BDI | 7BDJ | 7BDK | 7BDL |

<sup>a</sup> Values in parentheses refer to the highest resolution shells.
<sup>b</sup> $R_{\text{meas}}(I) = \sum_h [N/(N-1)]^{1/2} \sum_i |I_{ih} - \langle I_h \rangle| / \sum_h \sum_i I_{ih}$ , in which $\langle I_h \rangle$ is the mean intensity of symmetry-equivalent reflections $h$ , $I_{ih}$ is the intensity of a particular observation of $h$ and $N$ is the number of redundant observations of reflection $h$ .
<sup>c</sup> $CC_{1/2} = (\langle I^2 \rangle - \langle I \rangle^2) / (\langle I^2 \rangle - \langle I \rangle^2) + \sigma_e^2$ , in which $\sigma_e^2$ is the mean error within a half-dataset (84).
<sup>d</sup> $R_{\text{work}} = \sum_h |F_o - F_c| / \sum F_o$ (working set, no $\sigma$ cut-off applied).
<sup>e</sup> $R_{\text{free}}$ is the same as $R_{\text{work}}$ , but calculated on the test set of reflections excluded from refinement.
<sup>f</sup> A.U. – asymmetric unit.
<sup>g</sup> Calculated with Phenix.
<sup>h</sup> Rmsd - root-mean-square deviation from target geometry.

**Table S2.** Number of hydrogen bonds between protein and nucleotides.

| Cassette | NC-CC<sup>ATP</sup> | NC<sup>ADP</sup>-CC | NC<sup>ADP</sup>-CC<sup>ATP</sup> | NC<sup>ATP</sup>-CC<sup>ATP</sup> |
| --- | --- | --- | --- | --- |
| NC |  | 7.8±0.8 | 7.0±1.3 | 7.5±2.7 |
| CC | 4.0 ±0.9 |  | 7.1±1.1 | 6.8±1.1 |

**Table S3.**
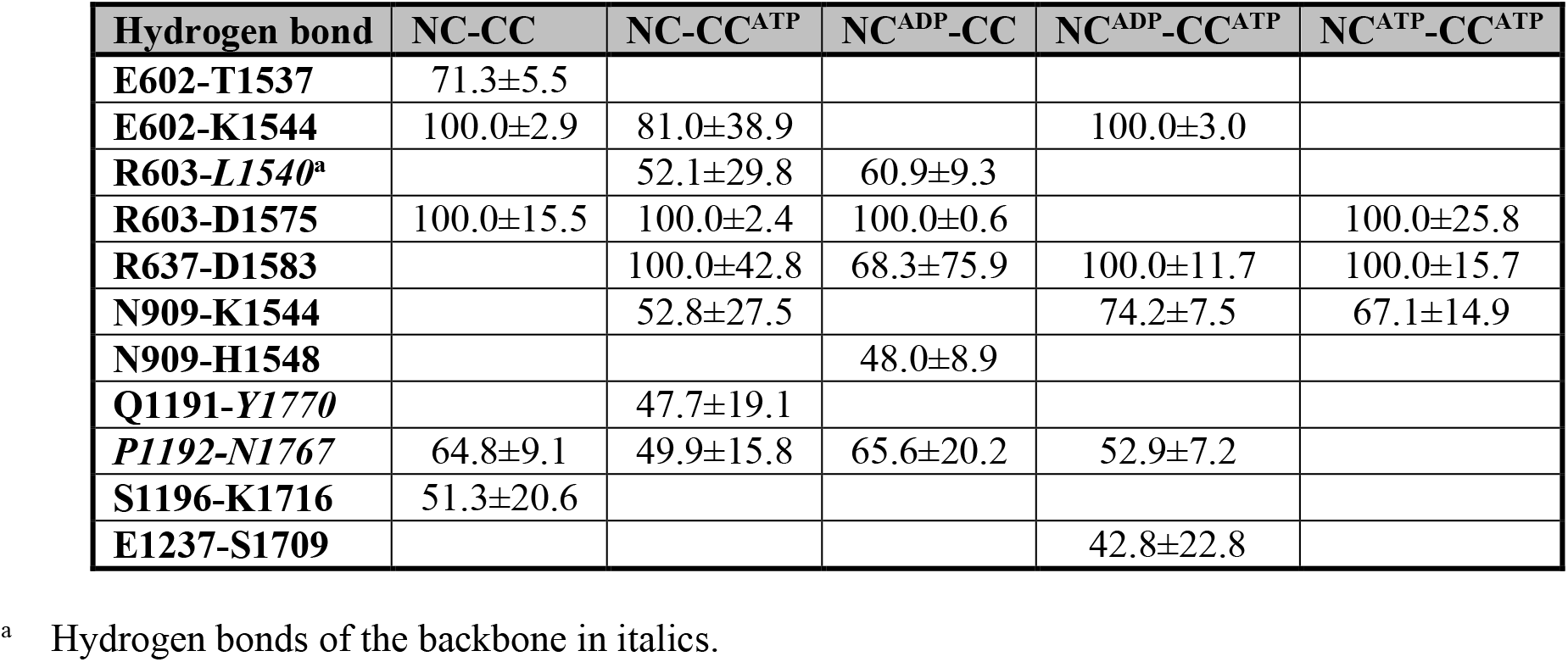
Hydrogen bond occupancies between the two cassettes [%].

**Table S4.** Crossing of cassette-cassette interface on shortest paths.

| Path | NC-CC | NC-CC<sup>ATP</sup> | NC<sup>ADP</sup>-CC | NC<sup>ADP</sup>-CC<sup>ATP</sup> | NC<sup>ATP</sup>-CC<sup>ATP</sup> |
| --- | --- | --- | --- | --- | --- |
| **E616-E1455** | R637-D1583 | E602-K1544 | R637-D1583 | R603-L1540 | E602-K1544 |
| **K509-K1356** | R637-D1583 | E602-K1544 | Y605-H1534 | R603-D1575 | Q980-Q1528 |
| **N820-N1692** | K599-H1534 | E602-K1544 | Y605-H1534 | R603-L1540 | E602-K1544 |
| **Q484-Q1332** | E578-H1534 | E602-K1544 | Y605-H1534 | R603-D1575 | E602-K1544 |
| **R855-E1455** | R637-D1583 | E602-K1544 | R637-D1583 | R603-L1540 | E578-H1534 |
| **T510-T1357** | R637-D1583 | M641-T1578 | Y605-H1534 | R603-D1575 | E602-K1544 |

## SUPPLEMENTAL FIGURES

**Figure S1.**
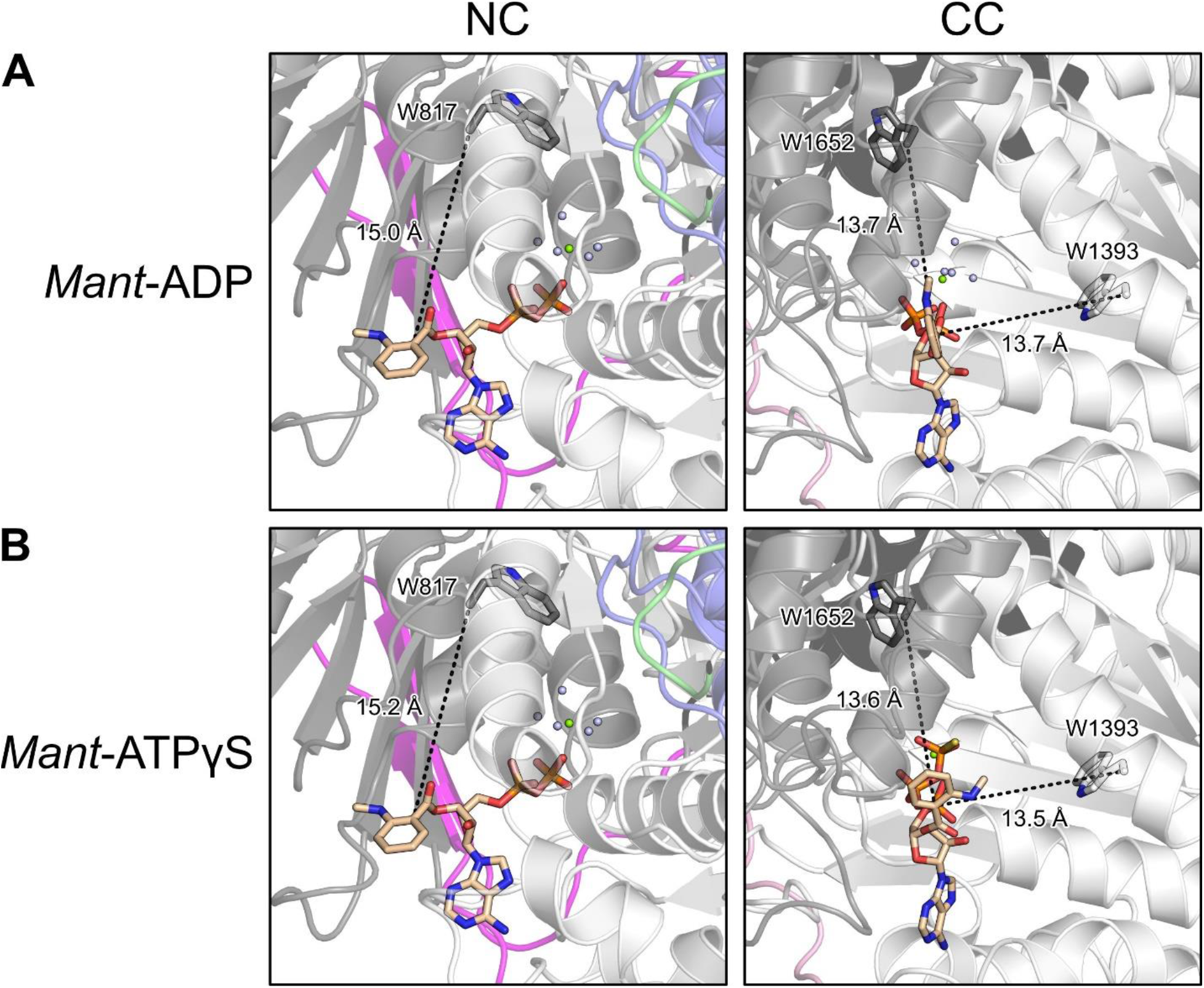
Distances of Trp residues to *mant* moieties. **A** and **B**. Comparison of the distances of the *mant* moieties in *mant*-ADP (**A**) or *mant*-ATPγS (**B**) to the nearest Trp residues around the NC (left) and CC (right) nucleotide binding pockets. Dashed lines, distances between the C8 atom of the *mant*-nucleotides to the Cα atoms of the respective Trp residues.

**Figure S2.**
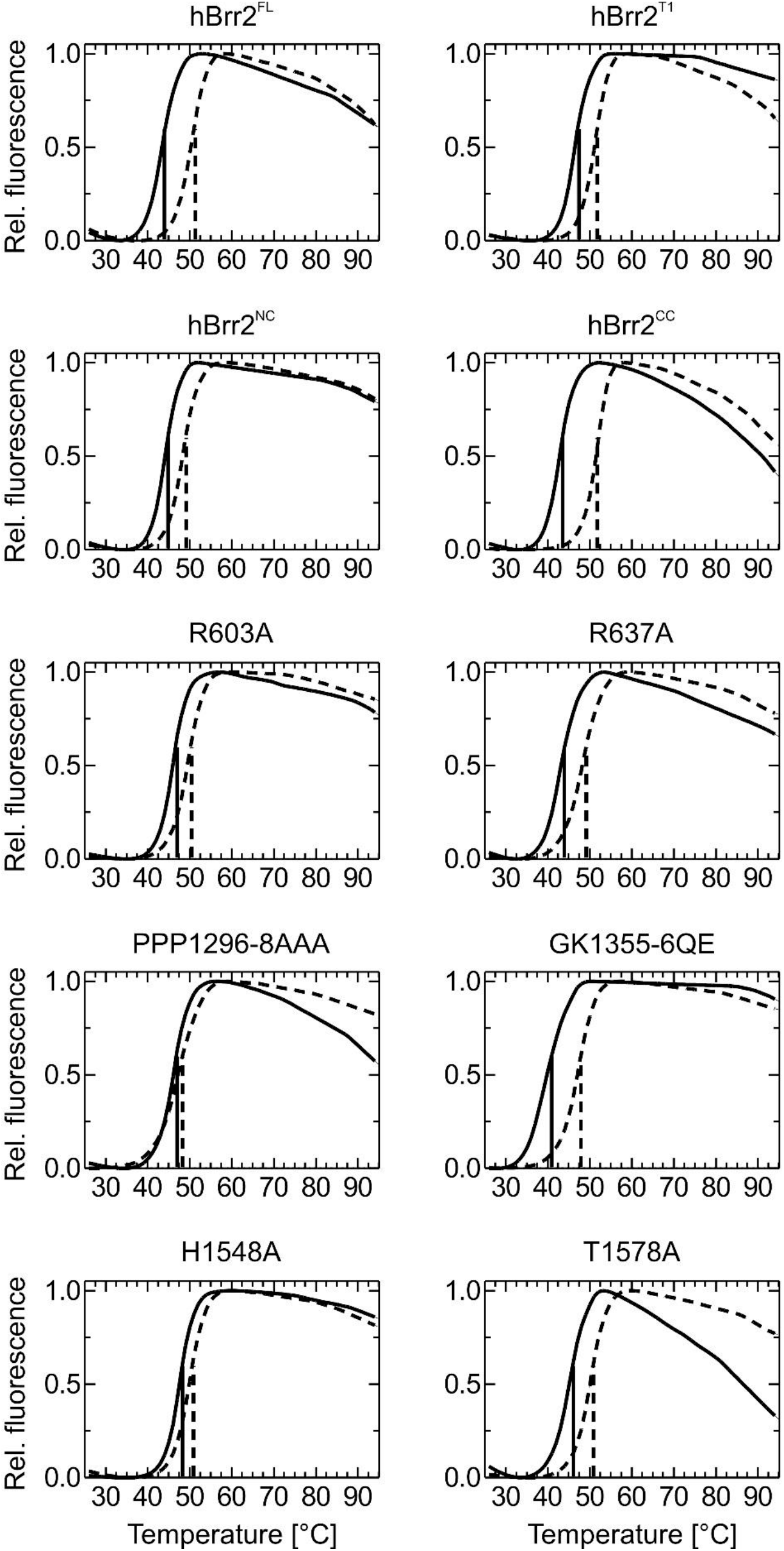
DSF analyses of the hBrr2 variants used in this work. Solid lines; melting curves obtained in the absence of ATP/Mg^+2^; dashed lines, melting curves obtained in the presence of 2 mM ATP/Mg^+2^. All variants exhibited cooperative transitions with similar melting temperatures.

